# B cells maintain the homeostasis of splenic marginal zone antigen-presenting cells to promote the anti-viral CD8^+^ T cell response

**DOI:** 10.1101/2025.04.28.651045

**Authors:** Xinyuan Liu, Filiz Demircik, Mariia Antipova, Stylianakis Emmanouil, Matthias Klein, David Bejarano, Abdelrahman Elwy, Anna Ebering, Michaela Blanfeld, Katlynn Carter, Lisa Johann, David Uhlfelder, Elisa Blickberndt, Hans Christian Probst, Nadine Hövelmeyer, Tobias Bopp, Ramon Arens, Joke M.M. den Haan, Jennifer L Gommerman, Esther von Stebut, Björn E Clausen, Andreas Schlitzer, Karl S Lang, Ronald A Backer, Niels A Lemmermann, Ari Waisman

## Abstract

Natural killer and CD8^+^ T cells are critical in the elimination of blood-borne viruses such as cytomegalovirus (CMV); however, the role of B cells in this process is less clear. Here, using the murine CMV (MCMV) infection model, we demonstrated that the B cell-deficient mice mounted a weaker primary virus-specific CD8^+^ T cell response than their wild-type counterparts, which was associated with increased viral transcription. Notably, we found that the contribution of B cells to the CD8^+^ T-cell-mediated anti-viral response was not associated with their ability to generate antibodies but with their ability to sustain Langerin^+^ type 1 conventional dendritic cells (cDC1s), a dendritic cells (DC) subset known for being involved in viral and bacterial clearance in the marginal zone of the spleen. Furthermore, we found that the presence of Langerin^+^ cDC1s is dependent on B cells expressing lymphotoxin (LTβ) to maintain CD169^+^ marginal metallophilic macrophages (MMMs). We further discovered, using ligand-receptor interaction analyses, that the communication between MMMs and Langerin^+^ cDC1s was mediated via VCAM1 - ITGA4/ITGB1 interaction. Thus, our data reveals that B cell regulate the development of MMMs in the spleen via LTβ expression and consequently sustain Langerin^+^ cDC1s homeostasis for effective initiation of an anti-viral CD8^+^ T cell response. Overall, our study offers a new perspective on how B cells maintain the homeostasis of antigen-presenting cells in the splenic marginal zone and thus indirectly affect the virus-specific CD8^+^ T cell response, which could potentially be extended to other infectious and autoimmune diseases as well as tumors.

## INTRODUCTION

Cytomegalovirus (CMV), which belongs to the β subfamily of herpesviruses, establishes lifelong latent infection in humans (*1*). Because CMV is strictly species-specific, murine CMV (MCMV) serves as a well-established animal model for human CMV (HCMV) pathogenesis and immune control (*2, 3*). Although HCMV does not typically cause symptomatic disease in immunocompetent individuals, it is often reactivated in immunocompromised bone marrow (BM) transplant recipients, causing severe disease (*4, 5*). In these patients, clearance of CMV infection relies on the efficient reconstitution of the CD8^+^ T cell repertoire (*6, 7*). Similarly, in immunocompromised mice, the transfer of MCMV-specific CD8^+^ T cells plays a protective role (*3, 8, 9*). In contrast to T cells, the significance of B cells during acute CMV infection is less clear (*10*).

B cells play an important role in splenic architecture development, particularly the maintenance of the marginal zone (MZ) (*11*). The splenic MZ is a distinctive and specialized structure, which is located at the interface between the lymphoid white pulp and the scavenging red pulp of the spleen. On gaining entry into the body, blood-borne viruses such as CMV first interact with immune cells in the splenic MZ. Here, the interplay between innate and adaptive immune mechanisms ensures an early and efficient response to pathogen-associated antigens (*12, 13*). The splenic MZ plays an important role in the immune response to blood-borne viral infections because it is enriched with antigen-presenting cells (APCs), including CD169^+^ marginal metallophilic macrophages (MMMs), SIGN-R1^+^ MZ macrophages (MZMs) and conventional dendritic cells (cDCs). Collectively, these cells capture antigens from the circulation and process them for clearance or presentation to cells of the adaptive immune response (*14–16*). CD169^+^ macrophages are reported to be the primary targets of viral infection (*17–20*). CD169, expressed on the surface of MMMs, is involved in cell-cell contact and transfers viral antigens to CD8^+^ cDC1s, which in turn prime the anti-viral CD8^+^ T cell response (*21*). MMMs also promote viral replication, which can actually increase the delivery of viral antigens to T and B cells and strengthen the anti-viral immune response (*22, 23*). cDCs are commonly classified into type 1 and type 2 lineages (cDC1s and cDC2s, respectively) (*24*), which can be further subdivided into several unique cDC subsets with various immune-modulatory roles (*25–29*). One of these subsets are Langerin^+^ cDC1s, which are a professional cross-presenting subgroup of CD8^+^ cDC1s that express high levels of DEC205, Clec9a, and CD36 and can assimilate circulating apoptotic cell-associated antigens (*30*). In addition, upon systemic stimulation, Langerin^+^ cDC1s produce large amounts of IL-12, a cytokine that is important for the clearance of viral and bacterial infections (*31, 32*). However, our understanding of whether MMMs and Langerin^+^ cDC1s can modulate primary anti-viral CD8^+^ T cell responses is limited. Moreover, the relationship between B cells, MMMs, and Langerin^+^ cDC1s is unclear.

Here, we demonstrated that B cells were indirectly required for the effective initiation of the anti-viral CD8^+^ T cell response, mainly because absence of B cells leads to a reduction in the number of Langerin^+^XCR1^+^ cDC1s, which were essential for the priming of a robust MCMV-specific CD8^+^ T cell response. Moreover, we found that B cells are required to maintain MMMs in the first place by expressing lymphotoxin β (LTβ), which in turn maintains the homeostasis of Langerin^+^XCR1^+^ cDC1s in the splenic MZ. By unbiased ligand-receptor network analysis, we identified the dialogue between MMMs and Langerin^+^ cDC1s involving VCAM1 - ITGA4/ITGB1. Taken together, these results provide a mechanistic explanation for how B cells regulate the primary MCMV-specific CD8^+^ T cell response, which may be applicable to other disease models.

## RESULTS

### B cells enhance primary CD8^+^ T cell responses to MCMV infection

To explore the role of B cells in the primary CD8^+^ T cell response, we first infected control and B cell-deficient J_H_T mice with MCMV. After seven days, we isolated the splenocytes of the infected mice and stimulated them with previously reported antigenic MCMV peptides *in vitro* (*33*). The activity of CD8^+^ T cells was assessed by measuring their capacity to produce IFN-γ. We found that the MCMV-infected J_H_T mice had markedly reduced frequencies of IFN-γ-producing CD8^+^ T cells compared to the infected control mice (Fig. 1A). To determine whether the decrease of the primary CD8^+^ T cell response in J_H_T mice was due to the lack of antibodies, we also infected IgMi mice with MCMV. IgMi mice harbor a polyclonal population of B cells but are not able to class switch or to produce secreted antibodies (*34*). We found that the percentage of IFN-γ-producing CD8^+^ T cells in the IgMi mice was similar to that of controls (Fig. 1A). To directly examine the role of immunoglobulins in the acute MCMV-specific CD8^+^ T cell response, we treated J_H_T mice with normal mouse serum (NMS) or mouse serum from MCMV-infected wildtype (WT) mice (IMS). In agreement with the results of the IgMi infection experiment, transfer of serum from infected WT mice did not restore the virus-specific CD8^+^ T cell response during acute MCMV infection (Fig. 1B). Together, these data indicate that secreted antibodies do not play a role in the primary anti-CMV CD8^+^ T cell response. Next, we assessed whether the reduction in virus-specific activated CD8^+^ T cells affected viral replication. To this end, we isolated the lymph nodes (LNs) of the MCMV-infected control and J_H_T mice and quantified viral transcription. Indeed, the weakened virus-specific CD8^+^ T cell response in the J_H_T mice was accompanied by higher MCMV transcription (Fig. S1A). This finding suggests that early MCMV control is less effective in B cell-deficient than control mice.

**Fig. 1.**
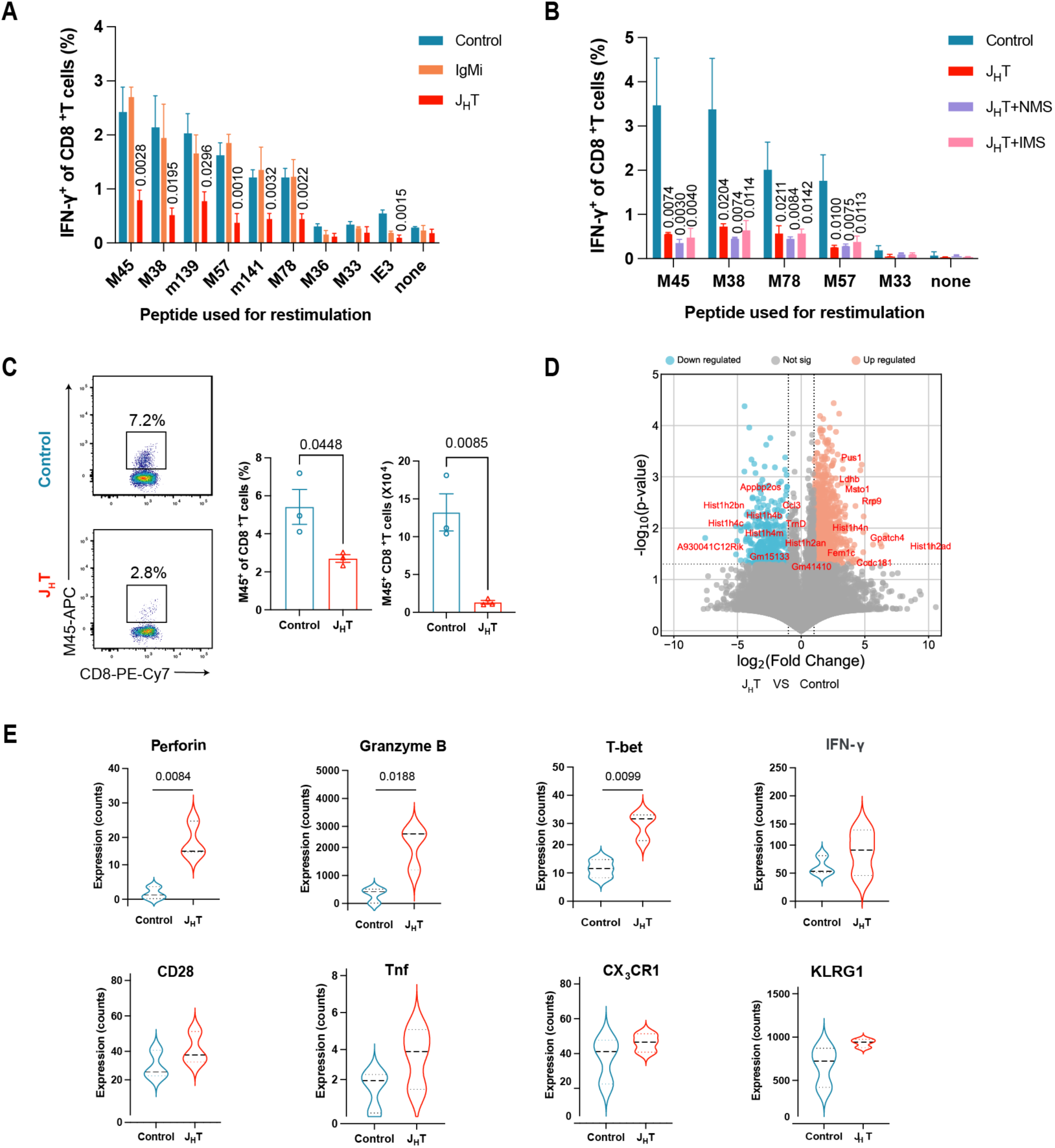
B cell deficiency attenuates the primary CD8^+^ T cell response to MCMV infection. (**A**) Frequency of IFN-γ-producing CD8^+^ T cells in the spleens after stimulation with MCMV peptides at 7 days post-infection of control, J_H_T, or IgMi mice (n = 3–12). (**B**) Frequency of IFN-γ-producing CD8^+^ T cells in the spleens of MCMV infected J_H_T and control mice after stimulation with MCMV peptides. Two of the conditions involved treating the J_H_T mice with normal mouse sera (NMS) or-infected mouse sera (IMS) before the MCMV infection (n = 3–4). (**C**) Representative flow cytometry dot plots of MCMV-M45 tetramer^+^ CD8^+^T cells in the spleens of MCMV infected control and J_H_T mice at 7 days post-infection. Bar graphs indicate the frequencies (left) and absolute numbers (right) of MCMV-M45 tetramer^+^ CD8^+^T cells in each group (n = 3). (**D**) Volcano plots depicting differentially expressed genes (DEGs) in splenic MCMV-M45 tetramer^+^ CD8^+^T cells of MCMV infected J_H_T versus MCMV infected control mice (P-value versus log2 fold change). The top 10 upregulated and downregulated DEGs are labelled. Genes with an adjusted P-value < 0.05 and log2 [fold change] > 1 or < −1 were considered as significantly differentially expressed. (**E**) Violin plots showing the expression of selected genes associated with the anti-viral activity of T cells in splenic MCMV-M45 tetramer^+^ CD8^+^T cells. Data are presented as the mean ± SEM and representative of two (1A, 1B,1C) independent experiments. Bulk RNA sequencing data were obtained from three control and three J_H_T mice. Statistical analysis: one-way ANOVA, Fig. 1A, 1B. Student’s t-test, Fig. 1C.

To determine whether the transfer of B cells would restore the magnitude of the primary CD8^+^ T cell response in the B cell-deficient mice, we reconstituted J_H_T mice with magnetically-sorted splenic WT B cells, as previously described (*35*). One week after transfer, the mice were infected with MCMV and the virus-specific CD8^+^ T cell response was analyzed at 7 days post-infection (Fig. S1B). Notably, adoptive B cell transfer partially restored the primary CD8^+^ T cell response of the J_H_T mice (Fig. S1C). To investigate if the inability of the CD8^+^ T cells to respond to the MCMV antigens was due to a defect caused by the constitutive lack of B cells from birth, we used our previously published iDTR mice (*36*) crossed to CD19-Cre (*37*) and treated them with diphtheria toxin (DT) to ablate the B cells. Indeed, the inducible depletion of B cells resulted in a dramatically reduced CD8^+^ T cell response to MCMV antigens, upon infection with the virus (Fig. S1, D and E). Together, these findings indicate that the attenuation of the MCMV-specific CD8^+^ T cell response correlates with the absence of B cells in these mice.

Next, we investigated how the lack of B cells affected the virus-specific CD8^+^ T cells in ways other than reducing their IFN-γ production. To this aim, we first analyzed CD8^+^T cell frequencies in J_H_T mice at steady state and after MCMV infection. We found that the frequency and number of CD8^+^ T were similar in the spleens of J_H_T and control mice at steady state and after MCMV infection (Fig. S1, F and G). Subsequently, we characterized CD8^+^ T cells recognizing the MCMV immunodominant M45 peptide using MHC tetramers in MCMV-infected control and J_H_T mice. Consistent with the reduction in IFN-γ-producing MCMV-specific CD8^+^ T cells, there were fewer M45 tetramer-positive CD8^+^ T cells in J_H_T than in control mice (Fig. 1C). To determine the molecular pathways implicated in the defective CD8^+^ T cell response during MCMV infection, we sorted equal numbers of these antigen-specific CD8^+^ T cells from each group of mice and subjected them to bulk RNA sequencing. We identified 1,018 differentially expressed genes between the M45-specific CD8^+^ T cells isolated from J_H_T and control mice (Fig. 1D). Nevertheless, the expression of genes encoding molecules associated with anti-viral activity, such as CD28, killer cell lectinlike receptor subfamily G member 1 (KLRG1), perforin, granzyme B, TNF, CX_3_CR1, and IFN-γ was similar or even increased in the MCMV-specific CD8^+^ cells of J_H_T versus those of control mice (Fig. 1E). This result indicates that CD8^+^ T cells in J_H_T mice can become fully efficient effector CD8^+^ T cells after recognition of their cognate MCMV antigen. Thus, these results collectively show that the reduction in the magnitude of the primary CD8^+^ T cell response in J_H_T mice was not due to changes in the effector function of the CD8^+^ T cells themselves. Instead, the increased expression of effector molecules such as perforin and granzyme B by virus-specific CD8^+^ T cells of the J_H_T mice may be attributed to the higher viral load and the reduced number of virus-specific CD8^+^ T cells. In such a context, existing CD8^+^ T cells may boost their effector functions as a compensatory mechanism.

### Loss of B cells attenuates CD8^+^ T cell priming after viral infection

Changes in the function of APCs can impact the virus-specific primary CD8^+^ T cell response. To assess whether altered APC function was responsible for the reduction in MCMV-specific CD8^+^ T cell numbers in the J_H_T mice, we adoptively transferred Cell Trace-Violet-labeled CD8^+^CD45.1^+^ T cells isolated from OVA-specific TCR-transgenic OT-I mice into CD45.2^+^ J_H_T or control recipients. One day later, the mice were infected with recombinant MCMV encoding an immunodominant OVA peptide (MCMV-SIINFEKL) (*38*). At 3 days post-infection, we isolated splenocytes and analyzed the proliferation of the OT-I T cells expressing the Vα2Vβ5 OVA-specific TCR (Fig. S2A). In comparison to uninfected mice, infected control animals displayed a strong OT-I CD8^+^ T cell expansion, whereas transferred OT-I CD8^+^ T cells in J_H_T mice proliferated significantly less (Fig. 2A). In addition, the proportion and number of CD62L^−^CD44^+^ effector CD8^+^ T cells was significantly reduced among the OT-I T cells transferred into J_H_T mice compared to control mice (Fig. 2B). These results suggest that CD8^+^ T cells cannot be properly primed in the J_H_T mice, possibly due to a deficiency of functional APCs in these animals.

**Fig. 2.**
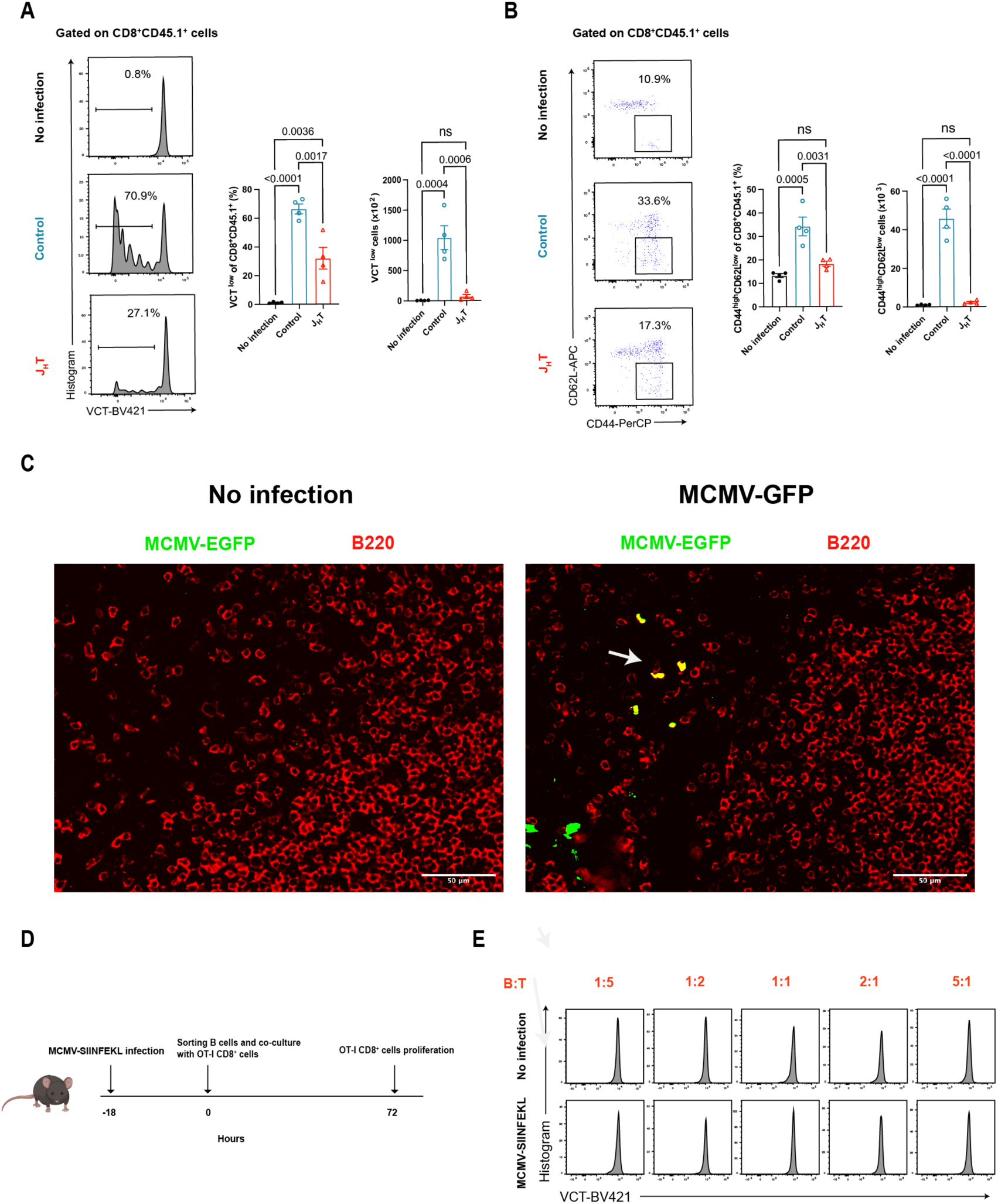
Absence of B cells impairs CD8^+^ T cell priming after viral infection (**A**) Proliferation of adoptively intravenous transferred OVA-specific transgenic OTI CD45.1^+^CD8^+^ T cells at 3 days after the induction of MCMV-SIINFEKL infection. Bar graphs indicate the frequencies (left) and absolute numbers (right) of proliferating OTI CD45.1^+^CD8^+^T cells in each group (n = 4). (**B**) Flow cytometry dot plot showing CD62L^-^CD44^+^ effector OTI CD45.1^+^CD8^+^ T cells at 3 days after the induction of MCMV-SIINFEKL infection. Bar graphs indicate the frequencies (left) and absolute numbers (right) of CD62L^-^CD44^+^ effector OTI CD45.1^+^CD8^+^T cells in each group (n = 4). (**C**) Immunofluorescence images of B220 and MCMV-EGFP in the spleens of 48 hours after MCMV-EGFP infection WT mice and uninfected control WT mice (scale bar, 50 µm; n = 4). (**D**) Schematic diagram outlining the program for co-culturing OVA-specific transgenic OTI CD8^+^ T cells with B cells isolated from MCMV-SIINFEKL Infected WT mice. (**E**) Flow cytometry histograms showing the proliferation of OVA-specific transgenic OTI CD8^+^ T cells co-cultured with B cells isolated from uninfected or MCMV-SIINFEKL-infected WT mice (n = 5). Data are presented as the mean ± SEM and representative of two independent experiments. Statistical analysis: One-way ANOVA, Fig. 2A, 2B.

B cells express both MHC class I and II molecules and can interact with T cells directly (*39, 40*). Thus, B cells can potentially act as APCs during the initiation of the MCMV CD8^+^ T cell response. In order to evaluate whether B cells are able to take up antigen after MCMV infection, and thus act as direct APCs, we infected WT mice with a recombinant MCMV expressing EGFP (MCMV-EGFP) (*41*). 48 hours later, splenic histology revealed a co-localization of B cells and MCMV-EGFP signals (Fig. 2C). Flow cytometry data also suggested that MCMV-GFP signals were detected in B cells (Fig. S2B). To investigate the ability of B cells to prime CD8^+^ T cells, we purified B cells from MCMV-SIINFEKL-infected WT mice and cultured them with OT-I T cells (Fig. 2D). We observed that the B cells isolated from these mice did not induce OT-I CD8^+^ T cell proliferation, suggesting that they were not the major APC type cross-presenting antigens to CD8^+^ T cells after MCMV infection (Fig. 2E). Together, these data prove that although the defective CD8^+^ T cell priming observed in the J_H_T mice was due to the lack of B cells, it did not rely on their ability to act as APCs to directly induce MCMV-specific CD8^+^ T cell responses.

### B cell deficiency affects the T cell priming capacity of XCR1^+^CD8^+^cDC1s

Having excluded the possibility that B cells acted as APCs to prime MCMV-specific CD8^+^ T cells directly, we presumed that the absence of B cells might affect other APCs central to this process, such as DC. To this end, we first investigated whether absence of B cells could change the ability of cDC1s to present MCMV Ags obtained in vivo. We infected control and J_H_T mice with MCMV-SIINFEKL, purified the XCR1^+^CD8^+^cDC1s from the spleens after 18 hours, and co-cultured them with CD8^+^ T cells from OT-I mice for 3 days at varying ratios (Fig. S3A). We found that co-culturing OT-I CD8^+^ T cells with XCR1^+^CD8^+^ cDC1s isolated from infected J_H_T but not control mice significantly reduced their proliferation (Fig. 3A). To further confirm that B cell deficiency affects the APC function of the CD8^+^ cDC1s, we isolated XCR1^+^ cDC1s from the spleens of naive J_H_T and control mice and incubated them with OVA protein. After 1 hour, the XCR1^+^cDC1s were washed and co-cultured with OT-I CD8^+^ T cells. After 3 days, we found that XCR1^+^ cDC1s from J_H_T mice expanded OT-I CD8^+^ T cells significantly less than those from control mice (Fig. 3B and Fig. S3B), demonstrating their inferior antigen-presenting capacity. In addition, the expression of co-stimulatory molecules CD40, CD80, and CD86 was significantly reduced in splenic cDC1s of J_H_T mice compared to control, consistent with the lower antigen-presenting function (Fig. S3C).

**Fig. 3.**
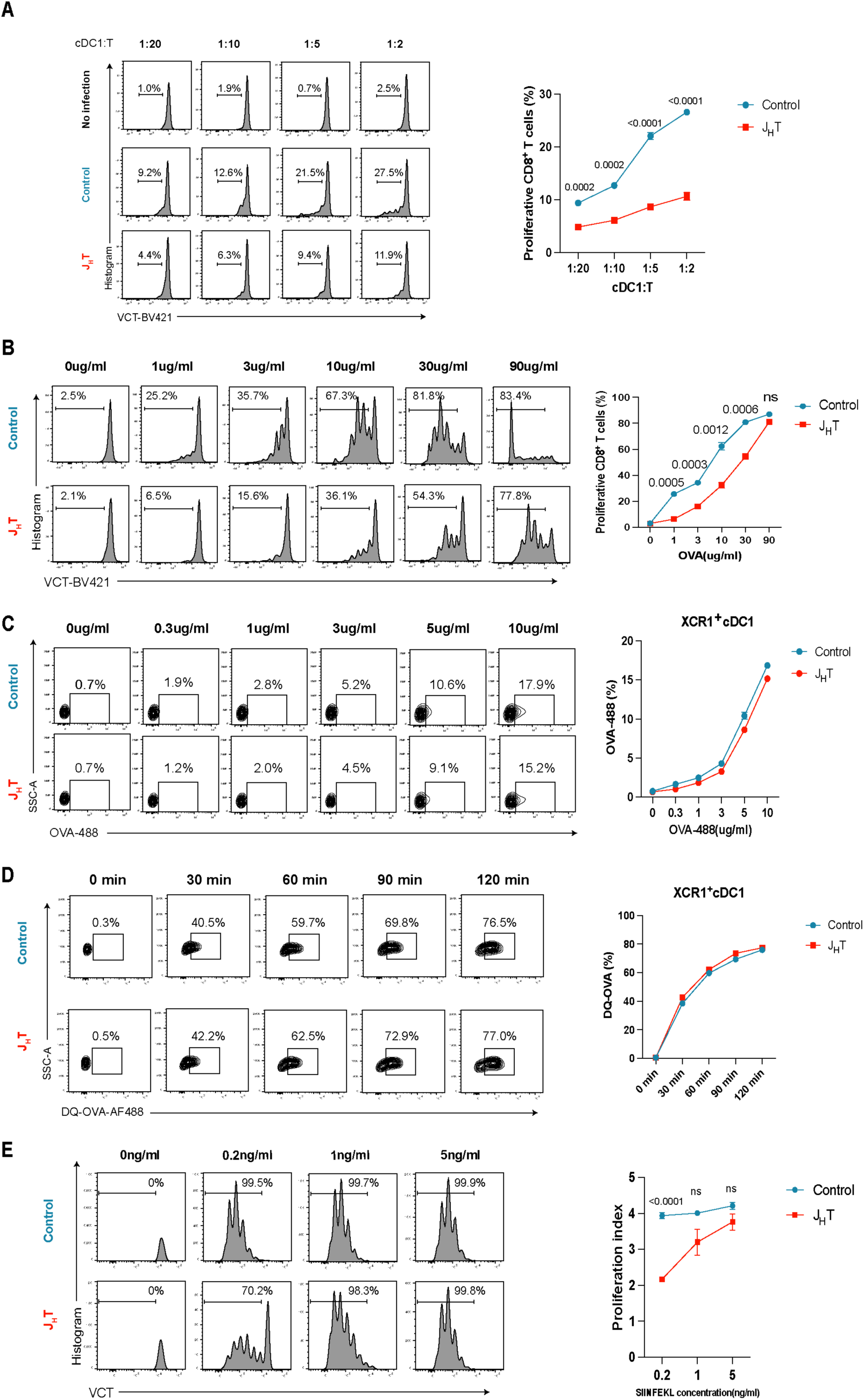
B cells support the CD8^+^-T-cell-priming function of XCR1^+^CD8^+^cDC1s. (**A**) Proliferation of OVA-specific transgenic OTI CD8^+^ T cells co-cultured at various ratios with cDC1s isolated from uninfected or MCMV-SIINFEKL-infected control and J_H_T mice. Line graph indicates the frequencies of proliferating OTI CD8^+^ T cells (n = 5). (**B**) Proliferation of OVA-specific transgenic OTI CD8^+^ T cells co-cultured with cDC1s isolated from control or J_H_T mice and pulsed with different concentration of OVA protein. Line graph indicates the frequencies of proliferating OTI CD8^+^T cells (n = 5). (**C**) Flow cytometry plots showing the frequencies of OVA-488^+^ XCR1^+^ cDC1s at each incubated concentration of OVA-488. Briefly, cDC1s isolated from control or J_H_T mice were incubated with increasing concentrations of Alexa-Fluor-488-labelled OVA for 30 minutes at 37 °C and 5% CO_2_. Line graph indicates the frequencies of OVA-488^+^ XCR1^+^ cDC1s in each group (n = 4). (**D**) Representative flow cytometry plots of DQ-OVA^+^ XCR1^+^ cDC1s exposed to DQ-OVA for different lengths of time. Briefly, cDC1s isolated from control or J_H_T mice were incubated with 10 μg/ml DQ-OVA for 0, 30, 60, 90, or 120 minutes at 37 °C and 5% CO_2_. Line graph indicates the frequencies of DQ-OVA ^+^ XCR1^+^ cDC1s in each group (n = 4). (**E**) Proliferation of OVA-specific transgenic OTI CD8^+^ T cells co-cultured with cDC1s isolated from control or J_H_T mice and pulsed with the SIINFEKL peptide. Line graph indicates the proliferation index of OTI CD8^+^T cells (n = 6). Data are presented as the mean ± SEM and representative of two (3A-3E) independent experiments. Statistical analysis: Student’s t-test, Fig. 3A-3E.

To better understand how the antigen-presenting function of XCR1^+^cDC1s was altered in J_H_T versus control mice, we evaluated their uptake of OVA conjugated to the Alexa Fluor 488 dye *in vitro*. We observed similar levels of green fluorescence in the XCR1^+^cDC1s of J_H_T and control mice, indicating that these cell populations exhibited similar antigen uptake (Fig. 3C). To determine whether the XCR1^+^cDC1s of J_H_T mice exhibited any antigen processing defects, which may have hampered MHC class I-mediated antigen presentation, we incubated XCR1^+^ cDC1s with OVA conjugated to the pH-insensitive BODIPY FL dye (DQ-OVA). After proteolytic degradation, the processed elements of DQ-OVA emit fluorescence, which can be analyzed by flow cytometry. We did not detect any differences in fluorescence between the groups, indicating that the splenic XCR1^+^ cDC1s of J_H_T and control mice were equally capable of processing phagocytosed DQ-OVA (Fig. 3D). Together, these data suggest that the cDC1s of B cell-deficient mice acquire and process antigens similarly to those of control mice. To examine other possible effects of B cell deletion on cDC1 function subsequent to antigen phagocytosis and processing, we co-cultured XCR1^+^ cDC1s and OT-I CD8^+^ cells in the presence of varying SIINFEKL peptide concentrations. We found that OT-I CD8^+^ T cells co-cultured with XCR1^+^ cDC1s from J_H_T mice expanded less at low SIINFEKL concentrations than those cultured with XCR1^+^ cDC1s from control mice (Fig. 3E). In accordance, the number of proliferating OT-I CD8^+^ T cells was significantly lower in co-cultures with XCR1^+^ cDC1s from J_H_T mice than in those with control XCR1^+^ cDC1s (Fig. S3C). Taken together, these data indicate that B cells contribute to the ability of splenic XCR1^+^ cDC1s to present antigen to CD8^+^ T cells.

### Loss of B cells reduces the number of the Langerin^+^XCR1^+^ cDC1 subset essential to prime the MCMV-specific CD8^+^ T cell response

Having shown that B cell deficiency reduced the CD8^+^ T cell priming capacity of cDC1s, we next explored the underlying mechanisms. First, we found that the lack of B cells did not affect the frequencies of cDCs in the spleen (Fig. S4A). Furthermore, there was no significant difference in the frequencies of SIRP-α^+^ cDC2 and XCR1^+^ cDC1 subsets in the spleens of J_H_T and control mice (Fig. 4A). We then employed high-dimensional flow cytometry to better characterize cDC subsets in the spleen, aiming to understand how they might be influenced by the B cell deficiency. Subsequently, we used t-distributed stochastic neighbor embedding (t-SNE) to show that the composition of splenic cDC1 and cDC2 populations differed significantly between J_H_T and control mice (Fig. 4B). Further characterization of the cDC1 and cDC2 subsets (Fig. S4B) revealed that the most significantly altered splenic cDC1 and cDC2 subpopulations in the J_H_T mice were Langerin^+^XCR1^+^ cDC1s and endothelial cell specific adhesion marker (ESAM)^hi^ SIRP-α^+^ cDC2s, respectively. In line with the t-SNE data, flow cytometry analysis of cDC1s or cDC2s revealed that the frequencies and numbers of Langerin^+^XCR1^+^ cDC1s and ESAM^hi^ SIRP-α^+^ cDC2s in the J_H_T mice were significantly lower than in control mice (Fig. 4C and Fig. S4C). In accordance, adoptive transfer of B cells into J_H_T mice partially rescued the Langerin^+^XCR1^+^ cDC1 subpopulation (Fig. S4D), as it restored the MCMV-specific CD8^+^ T cell response.

**Fig. 4.**
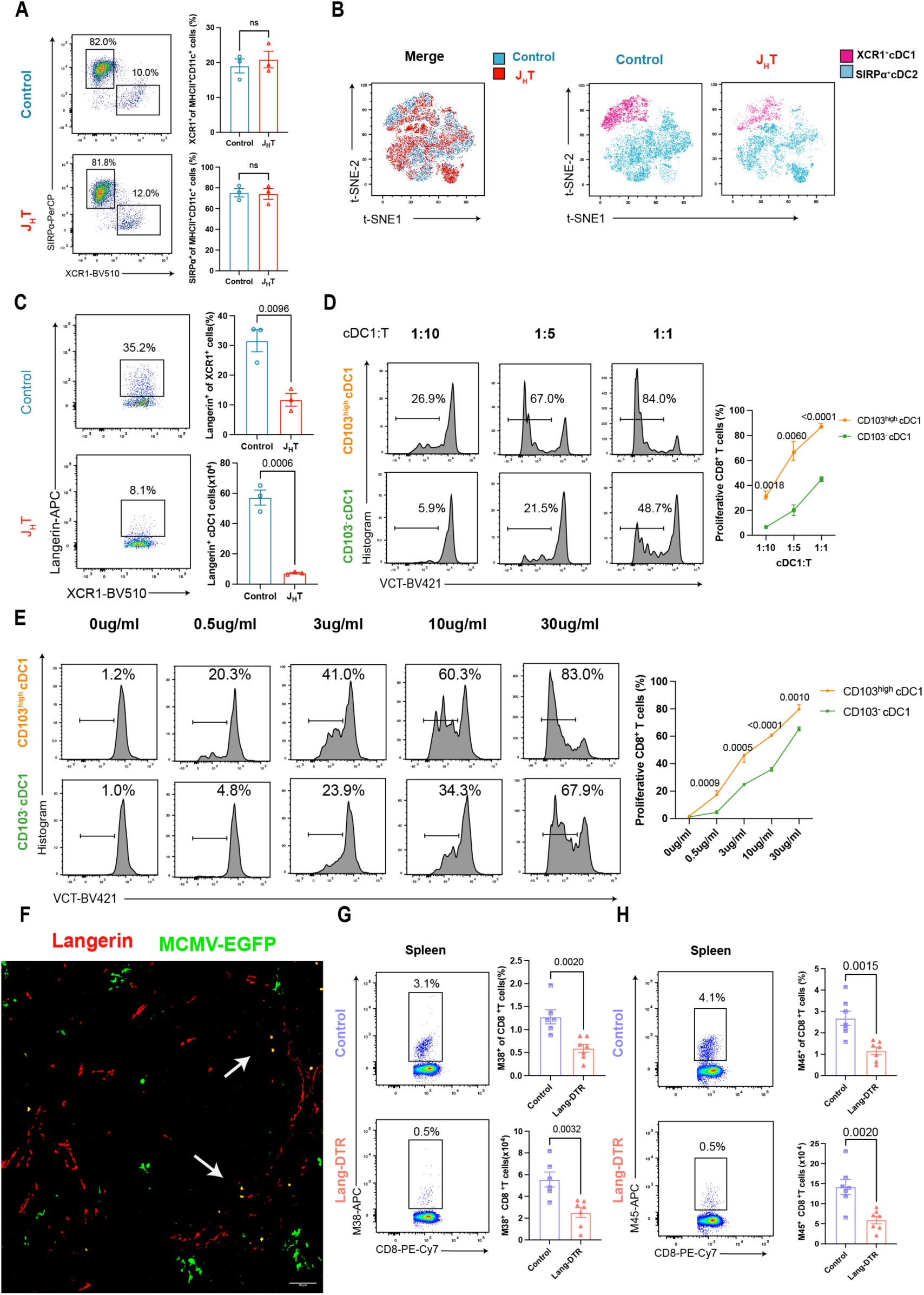
B cells maintain the homeostasis of splenic Langerin^+^ cDC1s. (**A**) Flow cytometry dot plot of XCR1^+^ cDC1s and SIRPα^+^ cDC2s subsets in the spleens of control or J_H_T mice. Bar graphs indicate the frequencies of XCR1^+^ cDC1s (top) or SIRPα^+^ cDC2s (bottom) (n = 3). (**B**) Three control and three J_H_T concatenated samples, each containing 5,000 splenic cDCs (CD11c^+^MHCII^+^CD19^-^), were embedded using the t-distributed stochastic neighbour embedding (t-SNE); cDC1s (pink) and cDC2s (blue) were classified according to XCR1 and SIRPα expression (n = 3). (**C**) Flow cytometry dot plot of splenic Langerin^+^ cDC1s from control or J_H_T mice. Bar graphs indicate the frequencies (top) and absolute numbers (bottom) of Langerin^+^ cDC1s (n = 3). (**D**) Proliferation of OVA-specific transgenic OTI CD8^+^ T cells co-cultured at various ratios with CD103^+^cDC1s or CD103^-^cDC1s isolated from MCMV-SIINFEKL-infected WT mice. Line graph indicates the frequencies of proliferating OTI CD8^+^ T cells (n = 9). (**E**) Proliferation of OVA-specific transgenic OTI CD8^+^ T cells co-cultured with CD103^+^cDC1s or CD103^-^cDC1s isolated from WT mice and pulsed with different concentration of OVA protein. Line graph indicates the frequencies of proliferating OTI CD8^+^T cells (n = 8). (**F**) Immunofluorescence images of Langerin and MCMV-EGFP in the spleens of WT mice 48 hours after MCMV-EGFP infection (scale bar, 50 µm; n = 4). (**G**) Representative flow cytometry dot plots of splenic MCMV-M38 tetramer^+^ CD8^+^T cells in MCMV infected control and MCMV infected Lang-DTR mice. Bar graphs indicate the frequency (top) and absolute numbers (bottom) of MCMV-M38 tetramer^+^ CD8^+^T cells (n = 7). (**H**) Flow cytometry plot of splenic MCMV-M45 tetramer^+^ CD8^+^T cells in MCMV infected control and MCMV infected Lang-DTR mice. Bar graphs indicate the frequencies (top) and absolute numbers (bottom) of MCMV-M45 tetramer^+^ CD8^+^T cells (n = 7). Data are presented as the mean ± SEM and representative of five (4A and 4C) or two (4B,4D-4G) independent experiments. Statistical analysis: Student’s t-test, Fig. 4A, 4C, 4D, 4E, 4G, and 4H.

Next, we asked whether the reduction in Langerin^+^cDC1s was responsible for the attenuated function of splenic XCR1^+^CD8^+^cDC1s to prime T cells in J_H_T mice. In splenic CD8^+^cDC1s, Langerin expression is mainly intracellular, and interestingly, the expression levels of the co-stimulatory molecules CD80 and CD86 are slightly higher in the Langerin^+^ cDC1 subset compared to Langerin^-^cDC1s at steady state (*30, 42, 43*), which may suggest that Langerin^+^cDC1s are functionally more mature. To assess whether Langerin^+^ cDC1s are efficient presenters of MCMV antigens they phagocytosed *in vivo*, we infected WT mice with MCMV-SIINFEKL, purified Langerin^+^cDC1s from the spleens 18 hr later, and co-cultured them with OT-I CD8^+^ T cells. Considering that Langerin expression in splenic cDC1s is mainly intracellular, and many studies have shown that Langerin^+^cDC1s co-express high levels of the integrin CD103 (*14, 42*), we used CD103 as a sorting marker for the Langerin^+^cDC1 subset. At 72 hr of co-culture, we observed that CD103^high^cDC1s more efficiently expanded OT-I CD8^+^ T cells, as compared to CD103^-^cDC1s (Fig. 4D). In order to investigate the antigen-presenting function of Langerin^+^cDC1 in more detail, we sorted CD103^high^cDC1s and CD103^-^cDC1s and cultured them with DQ-OVA. We found that the processed elements of OVA-DQ emit fluorescence similarly in CD103^high^cDC1s and CD103^-^cDC1s (Fig. S4E), indicating that Langerin^+^cDC1s and Langerin^-^cDC1s excert a comparable ability to acquire and process antigen. Next, we isolated CD103^high^cDC1s and CD103^-^cDC1s from the spleens of WT mice and co-cultured them with OT-I CD8^+^ T cells *in vitro* in the presence of OVA protein. With this experimental setup, we could readily detect that CD103^high^cDC1s were able to induce more proliferation of CD8^+^T cells as compared to CD103^-^cDC1s (Fig. 4E). Together, these results suggest that Langerin^+^cDC1s exhibit a better antigen-presenting ability than Langerin^-^cDC1s and that the diminished CD8^+^T cell priming function of cDC1s in the spleens of J_H_T mice is due to a reduction in the Langerin^+^cDC1 subset.

Given the crucial role of Langerin^+^cDC1s in antigen presentation, we focused on how they contribute in MCMV infection. By spleen histology of WT mice infected with MCMV-EGFP, we found that Langerin staining co-localized with MCMV-EGFP signals (Fig. 4F). To determine if splenic Langerin^+^XCR1^+^ cDC1s were required for the control of MCMV infection, we treated Lang-DTR (*44*) and control mice with DT and subsequently infected them with MCMV. Multiple DT injections were well tolerated(*45*) and effectively depleted Langerin^+^XCR1^+^ cDC1s in the spleen (Fig. S4F). At 7 days post-infection, we assessed the virus-specific CD8^+^ T cell response by MCMV tetramer staining. The frequency and number of MCMV-specific tetramer-positive CD8^+^ T cells were significantly lower in the Lang-DTR mice than in the control mice (Fig. 4, G and H, and Fig. S4G). This finding indicates that the reduction in Langerin^+^XCR1^+^ cDC1s was responsible for the decrease in MCMV-specific CD8^+^ T cells observed in J_H_T mice. Collectively, our data suggest that the absence of B cells reduces splenic Langerin^+^XCR1^+^ cDC1 numbers, which may have altered the antigen-presenting function of the remaining cDC1s and ultimately hampers effective CD8^+^ T cell priming.

### B cell-expressed LTβ modulates CD169^+^ MMMs to maintain splenic Langerin^+^ cDC1s homeostasis

We next sought to investigate the mechanisms by which B cells sustain the homeostasis of splenic Langerin^+^cDC1s. Langerin^+^ cDC1s are predominantly localized in the MZ of the spleen(*14, 43*), and interestingly, it has been reported that B cells are able to support the development and maintenance of secondary lymphoid tissues such as the splenic MZ by expressing LTβ(*46, 47*). We therefore asked whether B cells could sustain the homeostasis of Langerin^+^cDC1s by expressing LTβ. To test this hypothesis, we generated mixed BM chimeras using BM from WT mice mixed with that of J_H_T mice (control) or using BM from LTβ-deficient mice mixed with that of J_H_T mice, generating mice in which only the B cells are unable to express LTβ. As expected, we found that the absence of B cell-expressed LTβ markedly reduced the Langerin^+^cDC1 population (Fig. 5A). Next, we infected the chimeric mice with MCMV and observed significantly fewer MCMV-tetramer-positive CD8^+^ T cells in the absence of B cell-expressed LTβ, mimicking the situation in B cell-deficient J_H_T mice (Fig. 5,B and C). Collectively, these data prove that the absence of B cell-expressed LTβ impairs CD8^+^ T cell priming in response to MCMV infection, likely due to inadequate Langerin^+^cDC1 numbers.

**Fig. 5.**
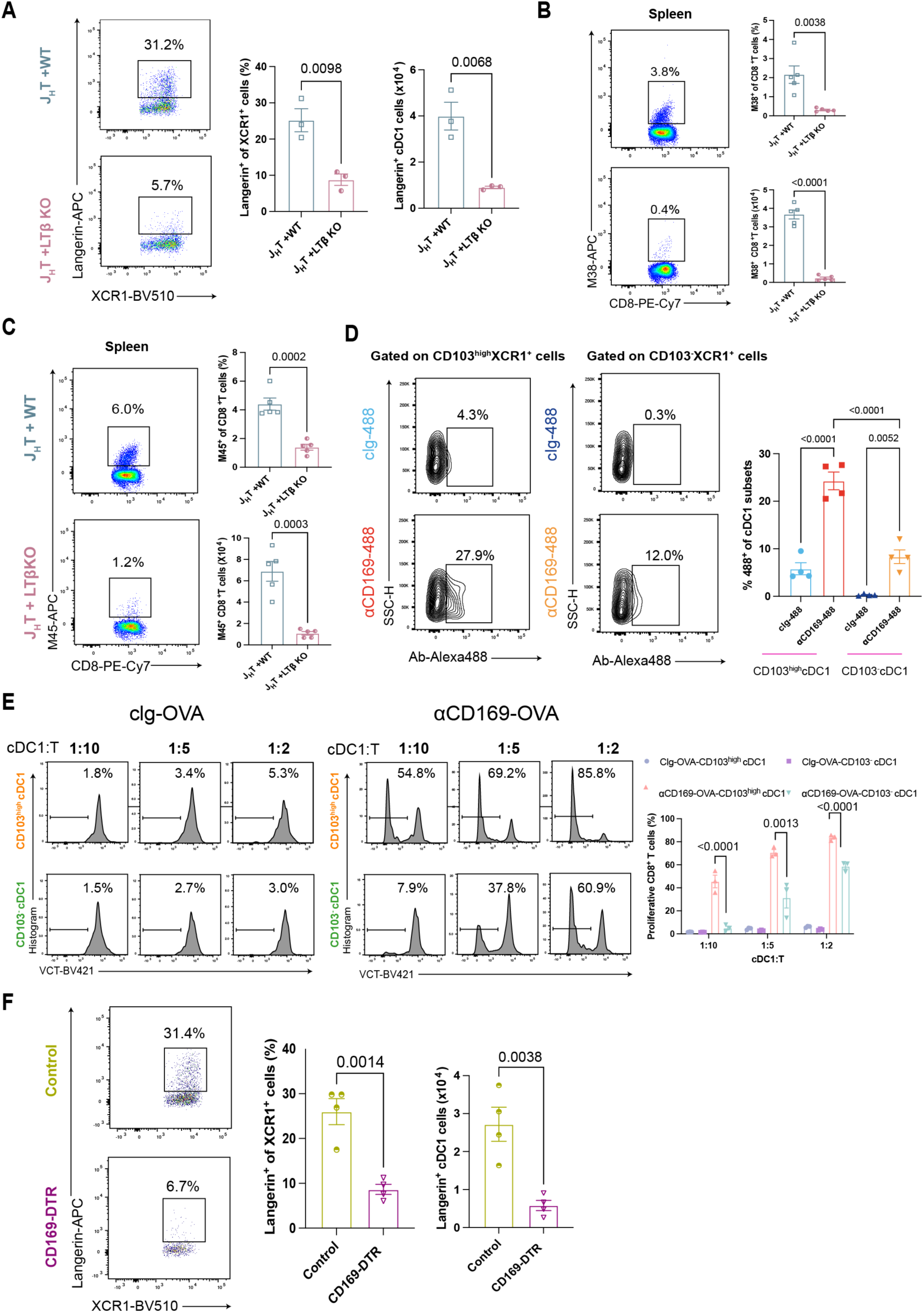
LTβ-expressing B cells maintain MMMs in the spleen to support the homeostasis of splenic Langerin^+^ cDC1s. (**A**) Flow cytometry plot showing splenic Langerin^+^ cDC1s in J_H_T mixed WT or J_H_T mixed LTβ-deficient BM chimera mice. Bar graphs indicate the frequencies (left) and absolute numbers (right) of Langerin^+^ cDC1s (n = 3). (**B**) Representative flow cytometry dot plots of splenic MCMV-M38 tetramer^+^ CD8^+^T cells in J_H_T mixed WT or J_H_T mixed LTβ-deficient BM chimera mice. Bar graphs indicate the frequencies (top) and absolute numbers (bottom) of MCMV-M38 tetramer^+^ CD8^+^T cells in each group (n = 5). (**C**) Flow cytometry dot plot showing splenic MCMV-M45 tetramer^+^ CD8^+^T cells in J_H_T mixed WT or J_H_T mixed LTβ-deficient BM chimera mice. Bar graphs indicate the frequencies (top) and absolute numbers (bottom) of MCMV-M45 tetramer^+^ CD8^+^T cells in each group (n = 5). (**D**) Flow cytometry plot showing Alexa488^+^ CD103^high^cDC1s and Alexa488^+^ CD103^-^cDC1s after indicated Ab-Alexa488 injection. Bar graphs indicate the frequencies of Alexa488^+^ CD103^+^cDC1s and Alexa488^+^ CD103^-^cDC1s (n = 4). (**E**) Proliferation of OVA-specific transgenic OTI CD8^+^ T cells co-cultured at various ratios with CD103^high^cDC1s or CD103^-^cDC1s isolated from indicated Ab-OVA immunized WT mice. Bar graphs indicate the frequencies of proliferating OTI CD8^+^ T cells. (**F**) Representative flow cytometry dot plots of splenic Langerin^+^ cDC1 cells from control or CD169-DTR mice. Bar graphs indicate the frequencies (left) and absolute numbers (right) of Langerin^+^ cDC1s (n = 4). Data are presented as the mean ± SEM and representative of two-three independent experiments. Statistical analysis: one-way ANOVA, Fig. 5D and 5E. Student’s t-test, Fig. 5A-5C, 5F.

Previous reports demonstrating that lymphotoxin (LT-α1β2)-LT-βR signaling regulates the development and/or homeostasis of splenic CD11b^+^ DCs, not cDC1s(*48–51*), and the absence of B cell-expressed LTβ also leads to alterations in the homeostasis of other cells in MZ(*47*), which led to wonder whether the regulation of Langerin^+^cDC1s by B cell-expressed LTβ needs to be dependent on other cells. As previously reported (*47, 52–54*), we also confirmed that lymphotoxin β (LTβ) expressed by B cells is required for the maintenance of MMMs within the splenic MZ (Fig. S5A). Interestingly, when we analyzed the spleens of WT mice at steady state, we found that Langerin^+^cDC1s colocalized with MMMs (Fig. S5B), suggesting that MMMs and Langerin^+^cDC1s could interact with each other. Previous studies reported that the functions of MMMs are Ag uptake and depend on CD169 binding to sialic acid on CD8^+^cDC1s to transfer Ag to CD8^+^ cDC1s for CD8^+^ T cell priming(*21, 55*). As langerin^+^cDC1s are a subset of CD8^+^ cDC1s, we hypothesized that MMMs can transfer antigen directly to them. To test this hypothesis, we injected WT mice with Alexa 488-labeled anti-CD169 (αCD169-488) antibody along with anti-CD40 and PolyI:C as adjuvants, and analyzed the transfer of αCD169-488 from MMMs to Langerin^+^cDC1s by fluorescence microscopy and flow cytometry. Indeed, we could confirm that MMMs are able to transfer antigen to Langerin^+^cDC1s (Fig. S5C). Moreover, we found that MMMs transferred more antigen to the CD103^high^cDC1 subset than to the CD103^-^cDC1 subset (Fig. 5D). Finally, we sorted CD103^high^ and CD103^-^ cDC1s from WT mice injected with αCD169-OVA and used them to stimulate OT-I CD8^+^T cells *in vitro*. In line with our previous results, OT-I CD8^+^T cell co-culture with CD103^high^cDC1s showed significantly increased proliferation compared to the CD103 negative counterpart (Fig. 5E), further supporting the concept that the Langerin^+^cDC1 subset is able to acquire more antigens from MMMs and present antigens more efficiently to CD8^+^ T cells.

Having established an intimate interaction between Langerin^+^ cDC1s and MMMs, our next objective was to determine whether B-cell-expressed LTβ regulated Langerin^+^XCR1^+^ cDC1s homeostasis is dependent on MMMs. Cytofluorometric analysis of DT-treated CD169-DTR mice revealed that MMMs depletion significantly decreased the frequencies and numbers of Langerin^+^XCR1^+^ cDC1s (Fig. 5D), but did not alter the presence of B cells (Fig. S5D,5E), indicating that MMMs play an essential role in maintaining splenic Langerin^+^XCR1^+^ cDC1s homeostasis and the regulation of Langerin^+^XCR1^+^ cDC1s homeostasis by B cells is dependent on MMMs. Moreover, the DT-treated Lang-DTR mice showed that Langerin^+^ cDC1s depletion did not affect the population of MMMs (Fig. S5F). Collectively, these data suggest that the lack of B cell-expressed LTβ disrupts the architecture of the spleen, and in particular, the localization of MMMs in the MZ, which in turn leads to a reduction in Langerin^+^XCR1^+^ cDC1s. Of note, here we cannot exclude the possibility that B cells also have the potential to regulate the homeostasis of Langerin^+^cDC1s directly through the expression of LTβ, but this study is difficult to perform because the inability of B cells to express LTβ leads to a significant reduction in MMMs, which in turn disrupts Langerin^+^cDC1s homeostasis.

As in the B cell-deficient mice, the antigen processing (Fig. S5G) abilities of cDC1s were not affected by the depletion of MMMs. Nevertheless, when XCR1^+^ cDC1s from the spleens of naive DT-treated CD169-DTR or control mice were loaded with OVA protein and then co-cultured with OT-I CD8^+^ T cells, as in the J_H_T mouse experiments, MMM depletion reduced the ability of XCR1^+^ cDC1s to prime OVA-specific OT-I CD8^+^ T cells (Fig. S5H). Taken together, these data suggest that the depletion of CD169^+^ MMMs perturbs the homeostasis of Langerin^+^XCR1^+^ cDC1s and affects the function of XCR1^+^ cDC1s to prime CD8^+^ T cells. These data clearly demonstrate that B cells maintaining splenic Langerin^+^ cDC1s homeostasis are dependent on self-expression of LTβ to sustain MMMs.

### VCAM1 - ITGA4/ITGB1 interaction coordinates MMMs - Langerin^+^XCR1^+^ cDC1s cross-talk

Next, we asked how MMMs maintain splenic Langerin^+^cDC1s homeostasis. For this purpose, we sorted CD11c^+^spleen cells from WT mice and performed single-cell sequencing (scRNA-Seq). Based on published splenic CD11c^+^ cell signatures and previously defined markers, we could define cDC1s, cDC2s, plasmacytoid DCs (pDCs), monocytes, and macrophages in our collected dataset (Fig. S6A). Subsequently, we selected MMMs and Langerin^+^cDC1s (Fig. S6B) and inferred cellular cross-talk based on ligand-receptor interactions between MMMs and Langerin^+^cDC1s from the analysis of scRNA-seq data. Individual ligand-receptor pair analysis suggested that VCAM-1-ITGA4/ITGB1 was the top-ranking ligand pair between MMMs and Langerin^+^cDC1s in WT mice (Fig. 6 A and B). Accordingly, VCAM1 and ITGA4 were respectively highly expressed in MMMs and Langerin^+^cDC1 clusters (Fig. 6C and Fig. S6C). Moreover, we have observed that CD169, Langerin, and ITGA4 co-localized in the spleens of WT mice as determined by fluorescence microscopy (Fig. 6D). To better understand whether MMMs could regulate the homeostasis of Langerin^+^cDC1s via VCAM1-ITGA4/ITGB1 interaction, we treated WT mice with the blocking anti-VCAM-1 antibody. While we did not observe any changes in the percentage and number of MMMs after anti-VCAM1 antibody treatment (Fig. S6D), but indeed VCAM-1 expression on MMMs was significantly reduced (Fig. S6E). Notably, our data indicate that the frequencies and numbers of splenic Langerin^+^XCR1^+^ cDC1s were significantly decreased after VCAM1 blockade (Fig. 6E). Next, we infected the anti-VCAM1 antibody treated mice with MCMV and assessed virus-specific CD8^+^ T cell response by MCMV tetramer staining at 7 days post-infection. We found the frequency and number of MCMV-specific tetramer-positive CD8^+^ T cells were significantly lower in the anti-VCAM1 antibody treatment mice than in the isotype treatment mice (Fig. 6F,G). Taken together, these results suggest that MMMs provide VCAM-1 signals, which are crucial for the maintenance of Langerin^+^XCR1^+^ cDC1s homeostasis in the spleen.

**Fig. 6.**
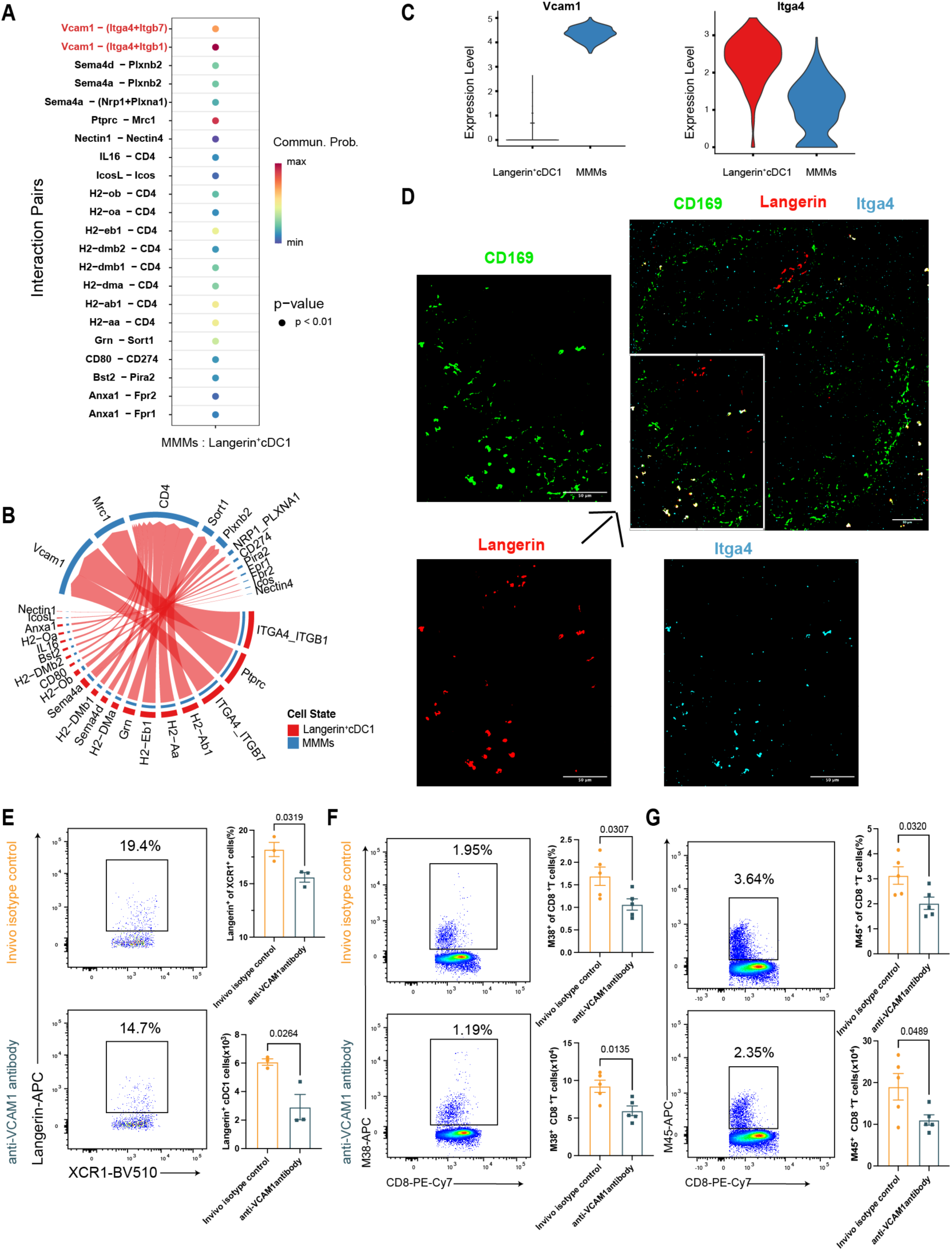
MMMs cross-talk with Langerin^+^XCR1^+^cDC1s by VCAM1 - ITGA4/ITGB1 **(A)** Ligand-receptor pair analysis between MMMs and Langerin^+^XCR1^+^cDC1s clusters. **(B)** Chord diagram showing preferential interactions between MMMs and Langerin^+^XCR1^+^cDC1 clusters. **(C)** Violin plots of VCAM1 and ITGA4 expression based on indicated cell subset. **(D)** Immunofluorescence images of CD169 (Siglec-1), Langerin, and ITGA4 in the spleens of WT mice (scale bar, 50 µm; n = 4). (**E)** Representative flow cytometry dot plots of splenic Langerin^+^ cDC1 cells from control IgG or anti-VCAM1 antibody treatment mice. Bar graphs indicate the frequencies (top) and absolute numbers (bottom) of Langerin^+^ cDC1s (n = 3). (**F)** Representative flow cytometry dot plots of splenic MCMV-M38 tetramer^+^ CD8^+^T cells in control IgG or anti-VCAM1 antibody treatment mice. Bar graphs indicate the frequencies (top) and absolute numbers (bottom) of MCMV-M38 tetramer^+^ CD8^+^T cells in each group (n = 5). (**G**) Flow cytometry dot plot showing splenic MCMV-M45 tetramer^+^ CD8^+^T cells in control IgG or anti-VCAM1 antibody treatment mice. Bar graphs indicate the frequencies (top) and absolute numbers (bottom) of MCMV-M45 tetramer^+^ CD8^+^T cells in each group (n = 5). Data are presented as the mean ± SEM and representative of two independent experiments (6D-6G). Statistical analysis: Student’s t-test, Fig. 6E-6G.

## DISCUSSION

CMV infection is a major reason for the high morbidity and mortality rates observed among immunologically compromised individuals, such as hematopoietic cell transplantation recipients and solid organ transplantation recipients; thus, CMV infection remains a compelling but under-addressed medical problem (*56, 57*). Our data reveal how B cells mechanistically contribute to the primary MCMV-specific CD8^+^ T cell response. In particular, we demonstrated that: 1) B cells play an important role in maintaining the homeostasis of Langerin^+^XCR1^+^ cDC1s, which are critical for efficient initiation of antiviral CD8^+^ T cell response; 2) The regulation of Langerin^+^XCR1^+^cDC1s homeostasis by B cells depends on their own expression of LTβ to maintain MMMs first, and MMMs in turn regulate the homeostasis of Langerin^+^XCR1^+^ cDC1s through the VCAM1 - ITGA4/ITGB1 cross-talk.

Besides producing antibodies, B cells modulate immune responses in a variety of ways, for example by producing pro-inflammatory cytokines such as TNFα and IL-6 or anti-inflammatory cytokines, including IL-10, IL-35, and TGF-β (*58–61*). These B cell-derived cytokines modulate the onset and development of autoimmunity in both mice and humans (*62–64*). Moreover, Graalmann et al. reported that B cells modulate the CD8^+^ T cell response to influenza or Modified vaccinia Ankara (MVA) infection (*65*). Consistent with these reports, we demonstrate that the absence of B cells from birth reduces the magnitude of the primary CD8^+^ T cell response to MCMV infection, which can be overcome by the adoptive transfer of B cells. Our data also indicate that the decreased MCMV CD8^+^ T cell response is not due to the antibody-secreting function of B cells. Many studies reported that B cells can also serve as professional APCs for CD4^+^ T cell priming (*66–68*); however, although B cells can capture virus-like particles similarly to DCs, they cannot efficiently process exogenous virus-like particles for cross-presentation in association with MHC class I (*69*). In the present study, we also observed that B cells do not effectively present antigens to naïve CD8^+^ T cells following MCMV infection, strongly suggesting that B cells modulate the MCMV-specific CD8^+^ T cell response via mechanisms other than direct antigen presentation.

B cells play crucial roles in the formation and maintenance of the splenic architecture, especially the MZ. The splenic MZ is enriched in APCs, including MMMs and Langerin^+^ cDC1s. The Langerin^+^ subset among cDC1s is particularly important for cDC1s functions such as cross-presentation and CD4^+^ Th1 cell stimulation (*14, 30, 42*). Our studies indicated that the abundance of Langerin^+^ cDC1s was significantly reduced in B cell-deficient mice. Prendergast et al. revealed that Langerin^+^ cDC1 play a pivotal role in the initiation of the CD8^+^ T cell response against bacteremia (*31*). Here, we used Lang-DTR mice to ablate Langerin^+^ cDC1s and in accordance with the above reports, we demonstrate that Langerin^+^ cDC1s played a critical role in the primary MCMV-specific CD8^+^ T cell response. Several reports suggest that MMMs can interact with cDC1s to promote anti-viral T cell responses (*21, 70, 71*). By immunizing mice with αCD169-488 antibody, we confirmed that MMMs could transfer antigen to Langerin^+^cDC1s, more importantly, our data suggest that MMMs are able to transfer antigens to CD103^+^ (Langerin^+^) cDC1s more efficiently. Interestingly, a recent study by Mauvais et al. found that MMMs process internalized antigens through an efficient vacuolar pathway and contribute to anti-tumor immunity independently of cDC1s(*72*). These studies all provide strong support that MMMs play an important role in antiviral CD8^+^ T-cell responses, and indeed, when we deleted MMMs by treating CD169-DTR mice with DT, we found that MCMV-specific CD8 T-cell responses were attenuated (data not shown). We further found that the presence of MMMs is essential for maintaining Langerin^+^ cDC1s homeostasis by characterizing the Langerin^+^ cDC1s of MMM-depleted mice. Thus, MMMs represent a direct link between B cells and Langerin^+^cDC1s, whereby B cell-expressed LTβ is a key factor in maintaining MMMs within the MZ (*47, 52, 53*), which is subsequently critical for the regulation of Langerin^+^cDC1s homeostasis. Through unbiased ligand-receptor network analysis, we demonstrated that MMMs shape Langerin^+^cDC1s homeostasis through the VCAM1-ITGA4/ITGB1 communication axis, but there may be other interactions between MMMs and Langerin^+^cDC1s that remain to be identified. Previous report has shown that loss of B cell-expressed LTβ also has altered marginal zone stromal cells(*47*), which are essential for MMMs development (*73*). Our study further proves that MMMs play an important role in regulating the homeostasis of Langerin^+^cDC1s. Altogether, our findings, together with previous studies, have provided a more detailed picture of how B cells regulate homeostasis of cells in the MZ.

Consistent with previous reports, our unsupervised visualization of high-dimensional flow cytometry data reveals that the frequency of ESAM^+^ cDC2s is reduced in B cell-deficient versus control mice(*50, 74*). Meanwhile, adoptive transfer of B cells into J_H_T mice partially rescued the ESAM^+^ cDC2 subpopulation (data not shown). cDC2s induce the activation of CD4^+^ T cells and their differentiation into Th2, Th17, and T-follicular helper cell subsets (*75–78*). Moreover, ESAM^+^ cDC2s, which are the major cDC2 subpopulation in the splenic MZ, are implicated in germinal center formation (*48, 79*). Therefore, utilizing different disease models to explore whether B cells can modulate CD4^+^ T cell responses through ESAM^+^ cDC2s represents an intriguing avenue for future research, which is currently hampered by the lack of cDC2-specific cell ablation mouse models.

Interestingly, B cell depletion is now considered one of the most effective treatments for multiple sclerosis (MS) (*80*), but the reason for the effectiveness of this therapy is not clear. Furthermore, some studies have shown that CD169^+^ macrophages are abundantly present in MS patients and promote neuroinflammation, suggesting that CD169^+^ macrophages play a key role in the pathophysiology of MS (*19, 81*). Based on our current findings and the fact that one can find clonal expansion of CD8^+^ T cells in the brains of MS patients (*82*), we suggest that the amelioration of disease seen in MS patients after B cell depletion may be due to reduced activation of autoreactive CD8^+^ T cells

In conclusion, our findings shed new light on the role of B cells in anti-viral immunity. Besides contributing to viral clearance through the production of antibodies and cytokines, our findings suggest that B cells critically contribute to the initiation of the virus-specific CD8^+^ T cell response by regulating other APCs such as macrophages and cDCs. These mechanistic insights will broaden our understanding of the multifaceted roles of B cells in shaping adaptive immune responses and offer new strategies for the treatment of immune-mediated diseases.

## MATERIALS AND METHODS

### Mice

All mice used were on the C57BL/6J background. The following mice have been previously described: J_H_T (*83*), IgMi (*34*), OT-I (*84*), C57BL/6J (*85*), CD45.1 (*86*), and bred in our facility (Translational Animal Research Center of the University Medical Center Mainz) under specific pathogen-free (SPF) conditions. For OT-I adoptive transfer experiments, we crossed OT-I mice with CD45.1^+^ mice to generate CD45.1^+^OT-I mice. CD169-DTR mice were provided by Lang KS. Lang DTR mice were obtained from Probst HC, and LTβ BM cells were provided by Gommerman JL. For all experiments, mice of both sexes were used at adult age, and the appropriate littermate mice were used as WT controls. All animal experiments were performed in accordance with the guidelines of the central animal facility institution.

### MCMV infection

Mice were infected via left hind footpad or with 1×10^5^ plaque-forming units (PFU) of cell-culture derived MCMV (strain Smith) or intravenously (i.v.) injection with the 2×10^5^ PFU of MCMV-SIINFEKL or i.v. with the 1×10^6^ PFU of MCMV-EGFP. Viral gene expression in the draining lymph node was quantified by RT-qPCR specific for m123/IE1 as described previously (*87*).

### Preparation of single cell suspensions and Cell Sorting

Cell suspensions for analysis of the DC population were purified from the spleen. Spleen was cut into grain size pieces and shaken for 30 minutes at 37°C in 1 ml of Roswell Park Memorial Institute (RPMI) 1640 media containing 200 U/ml Collagenase Type IV (Worthington Biochemical) and 0.5 U/ml DNaseI (Roche). After being digested, cell suspension was obtained by passing through a 70 μm cell strainer and washing in PBS with 10 mM EDTA and 2% Fetal Calf Serum (FCS), followed by centrifugation and lysis with ACK lysis buffer. To isolate MMMs, spleen was cut into small pieces in polypropylene tubes with 0.75 ml of RPMI that included 4 mg/ml Lidocaine (Sigma). In addition, 0.25 ml of RPMI containing 4 WU/ml of Liberase TL (Roche) and 2 U/ml of DNaseI (Roche) was added. Samples were then incubated at 37°C with constant stirring for 15 minutes until digested. After being digested, cells were passed through a 70 μm cell strainer, washed once with ice-cold RPMI plus 10% FCS, 20 mM Hepes, 10 mM EDTA, and 50 μM 2-mercaptoethanol, followed by centrifugation and lysis with ACK lysis buffer. To purify DC, we used anti-CD11c Magnetic-activated cell sorting (MACS) microbeads (Miltenyi Biotec) according to the manufacturer’s protocol. To sort XCR1^+^ CD8^+^ cDC1 populations, purified CD11c^+^ cells were stained for CD8a, CD11c, MHCII, XCR1, SIRPα, and viability dye. Subsequently, the ARIA III cell sorter (BD Biosciences) was used to sort cells based on the strong expression of CD11c and MHCII, as well as the presence of XCR1 and CD8a. After MACS or ARIA III sorting, the cells are reconfirmed to have >95% purity and viability.

### Flow Cytometry

Single cells were suspended in fluorescence-activated cell sorting (FACS) buffer (PBS/ 2mM EDTA/2% FCS) and blocked with anti-mouse CD16/32 (BioLegend) for 15min at 4°C. After blocking, cells were stained with the appropriate surface antibodies at 4°C in the dark for 20 minutes. MCMV-specific CD8^+^ T cells were stained with M38 or M45 tetramer at 4 °C in the dark for 40 min. Intracellular staining requires staining at 4°C for 1 hour or overnight after fixation and permeabilization using the FoxP3 staining kit (Invitrogen) according to the manufacturer’s instructions. According to the requirements of the experiment, we used the following antibodies for flow cytometry: anti-B220 (RA3-6B2), anti-CD4 (GK1.5), anti-CD8a (53-6.7), anti-CD11b (M1/70), anti-CD11c (N419), anti-CD19 (6D5), anti-CD44 (IM7), anti-CD45 (30-F11), anti-CD45.1 (A20), anti-CD62L (MEL-14), anti-CD103 (2E7), anti-CD106/VCAM1 (429(MVCAM.A)), anti-CD172/SIRP alpha (P84), anti-ESAM (1G8/ESAM), anti-F4/80 (BM8), anti-FceRIa (MAR-1), anti-IFN-γ (XMG1.2), anti-Ly-6C (HK1.4), anti-MHCII (M5/114.15.2), anti-TCR V alpha 2 (B20.1), anti-TCR Vbeta5.1 5.2 (MR9-4), anti-XCR1 (ZET) (all from Biolegend), and anti-CD64 (X54-5/7.1), anti-CD135 (A2F10.1), anti-Ly-6G (1A8) (all from BD Biosciences), and anti-CD169 (SER-4) (ThermoFisherScientific), anti-CD207/Langerin (122D5.03) (Eurobio Scientific). Flow cytometry data was required by FACS Canto II (BD Biosciences) or FACS Symphony (BD Biosciences) and analyzed using FlowJo v10 software.

### Immunofluorescence

Whole spleens were cut into 6-8 mm sections with a cryostat. Sections were dried overnight at room temperature, then stored at −80°C until staining began. Non-specific staining was blocked using PBS plus 5% normal goat serum and 0.5% Triton-100 for 1 hour at room temperature, then sections were stained at 4 °C overnight with the following primary antibodies: rat anti-CD169 (MOMA-1) (BMA BIOMEDICALS), rat anti-B220 (RA3-6B2) (BD Biosciences), rat anti-CD207/Langerin (eBioL31), and chicken anti-GFP (Novus). The next day, sections were washed and further labeled with the appropriate secondary antibodies (anti-rat IgG CF555 (Sigma Aldrich) or anti-chicken CF488A (Sigma Aldrich) for 1h at RT, followed by mounting with Fluoroshield mounting media with DAPI (Sigma-Aldrich). Images were captured by whole-slide scanning with a Zeiss SP8 and analyzed using Fiji software.

### Co-detection by indexing (CODEX) staining, imaging, processing and analysis

5µm slices of frozen spleens from control and MCMV-infected mice were prepared and used for CODEX staining as described before(*88*). Briefly, sections were dried for 15 min and fixed for 10 min in ice-cold acetone. Samples were subsequently rehydrated in hydration buffer and photobleached. Afterwards, sections were blocked and stained with Rat anti-human/mouse B220 (RA3-6B2) (Biolegend), Rat anti-mouse CD169 (3D6.112) (Biolegend), Rat anti-mouse CD207/Langerin (eBioRMUL.2) (Thermo Fisher Scientific), Rat anti-mouse CD49d (9C10) (Biolegend), and Rabbit anti-GFP (Thermo Fisher Scientific) antibodies overnight at 4°C. Samples were washed twice with staining buffer, fixed in ice-cold methanol, washed with 1x PBS, and fixed with BS3 fixative for 20min (Sigma Aldrich, St. Louis, MO, USA). After removal of the fixative, the stained samples were stored at 4°C before imaging. Then Coverslips and reporter plate were equilibrated at room temperature for 30min before imaging. A multicycle CODEX experiment was performed following manufacturer’s instructions (Akoya Biosciences). Images were acquired using a Zeiss Axio Observer widefield fluorescence microscope with a 20x objective (NA 0.85). A z-spacing of 1.5 µm was used for acquisition. The 405, 488, 568, and 647 nm channels were implemented. Raw files were exported using the CODEX Instrument Manager (Akoya Biosciences, Marlborough, MA, USA) and processed with CODEX Processor v1.7 (Akoya Biosciences). Images were inspected and analyzed in QuPath. After selecting the total cells using the DAPI counterstain, multiple classifiers were established using the single stains and used to gate the cells following a similar strategy as in flow cytometry.

### DC Phagocytosis assay

CD11c^+^ DC were isolated by CD11c MACS microbeads (Miltenyi Biotec) and diluted in medium to a concentration of 1.5 ×10^6^ cells/ml, and 100 μl of cells were plated in a 96-well round bottom plate. OVA-488 of the gradient concentration diluted with PBS (0 μg/ml, 0.3 μg/ml, 1 μg/ml, 3 μg/ml, 5 μg/ml, 10 μg/ml, 30 μg/ml, and 60 μg/ml) was added to the DC and incubated for 30 minutes at 37 °C in the cell culture incubator. The plate of an identical assay was kept on ice as a negative control. Excess fluorochrome bound to the cell surface was quenched by adding 0.1% Trypan blue stain and incubating for three minutes at room temperature. Afterwards, the cells were washed three times with PBS and then stained for flow cytometry.

### DC Processing assay

CD11c^+^ DC were isolated by CD11c MACS microbeads (Miltenyi Biotec) and diluted in medium to a concentration of 1.5 ×10^6^ cells/ml, and 100 μl of cells were plated in a 96-well round bottom plate. OVA-DQ was diluted in medium to a concentration of 2 μg/ml or 10 μg/ml. Diluted OVA-DQ was incubated with CD11c^+^ DC in a 37 °C cell culture incubator according to the following time points: 0 (OVA-DQ was added right before centrifugation), 30, 60, 90, and 120 minutes. The plate of an identical assay was kept on ice as a negative control. After incubation, the cells were centrifuged and stained for flow cytometry.

### Cell isolation and co-culture experiments

OVA-specific T cell transgenic (TCR) CD8^+^ T cell isolation was performed on the spleen and peripheral LNs from OT-I mice using the Mouse CD8a^+^ T Cell Isolation Kit (Miltenyi Biotec) according to the manufacturer’s instructions. For the cDC1/OVA or SIINFEKL/OT-I co-culture experiment, purified 4×10^4^ XCR1^+^ cDC1s or CD103^+^cDC1s or CD103^-^cDC1s was incubated with OVA or SIINFEKL peptide for 60 min at 37°C in the cell culture incubator; subsequently, XCR1^+^cDC1s or CD103^+^cDC1s or CD103^-^cDC1s were extensively washed and cultured with 1×10^5^ cell trace Cell Trace Violet (ThermoFisher) labeled OT-I CD8^+^ T cells. After 3 days, the proliferation of OT-I CD8^+^ T cells was analyzed by flow cytometry. In the MCMV-SIINFEKL infected cDC1s and OT-I CD8^+^ T cell co-culture experiment, purified XCR1^+^cDC1s or CD103^+^cDC1s or CD103^-^cDC1s from the spleen 18 hours after MCMV-SIINFEKL infection was cultured with cell trace violet (ThermoFisher) labeled OT-I CD8^+^ T cells with different ratios. The dilution of the cell dye was analyzed by flow cytometry on day 3 of culture. For the MCMV-SIINFEKL infected B cells and OT-I CD8^+^ T cell co-culture experiment, B cells were purified 18 hours after MCMV-SIINFEKL infection in WT mice using the MACS CD19 microbeads (Miltenyi Biotec) according to the manufacturer’s protocol. Purified B cells were cultured with cell trace violet (ThermoFisher) labeled OT-I CD8^+^ T cells with different ratios. After 3 days, the proliferation of OT-I CD8^+^ T cells was analyzed by flow cytometry.

### J_H_T mouse serum transfer

J_H_T mice were adoptively transferred with mouse serum derived from normal C57BL/6 mice and mouse serum derived from MCMV-infected C57BL/6J mice, 3 weeks after infection. The transfer started three days before infection, followed by one serum transfer on the day of MCMV infection and one serum transfer during the seven days of MCMV infection.

### B-cell transfer experiment

J_H_T mice were intravenously injected with 50×10^6^ cell MACS-purified B cells from the spleen of C57BL/6 mice (Miltenyi Biotec). One week after transfer, the recipients were infected in the footpads with 2×10^5^ PFU of MCMV. After seven-days post infection, splenocytes were analyzed for IFN-γ expression by *in vitro* stimulation with the MCMV peptides.

### OT-I adoptive transfer

Mice were i.v. injected with 5×10^6^ cell trace violet (ThermoFisher) labeled MACS purified CD8^+^T cells from the spleen and peripheral LNs of CD45.1^+^ OT-I mice (Miltenyi Biotec). Recipients were infected by intraperitoneal injection (I.P.) with 2×10^5^ PFU of MCMV-SIINFEKL after 24 hours. Spleens were removed 3 days after infection, and the proliferation of transferred CD45.1^+^ OTI CD8^+^T cells was analyzed using flow cytometry.

### In vivo Ag transfer Assay

Mice were immunized intravenously with 1 μg of αCD169-488 or clg-488 in the presence of 25 μg of poly(I:C) and 25 μg of αCD40. 2 hours after the injection, mice were sacrificed and spleens were frozen for immunofluorescence analysis or digested for (FACS) analysis.

### Ex Vivo Ag Presentation Assays

Mice were injected i.v. with 1 μg αCD169-OVA or clg-488 plus 25 μg αCD40 and 25 μg poly (I:C). 16 h later, purified CD103^+^cDC1s or CD103^-^cDC1s from the spleen was cultured with cell trace violet (ThermoFisher) labeled OT-I CD8^+^ T cells with different ratios. The dilution of the cell dye was analyzed by flow cytometry on day 3 of culture.

### Bone marrow chimeras

To generate mice in which B cells lacking LTβ expression, lethally irradiated C57BL/6J recipients were reconstituted with 80% J_H_T bone marrow plus 20% LTβ^−/−^bone marrow. Controls were created by reconstituting lethally irradiated C57BL/6J recipients with 80% J_H_T bone marrow plus 20% WT bone marrow. The recipient mice received antibiotic drinking water starting 10 days before being lethally irradiated and continuing for 6 weeks following bone marrow reconstitution. 8 weeks after reconstitution, mice were infected with MCMV and analyzed.

### VCAM1 blockade

Mice were injected i.v. with 10 mg/kg anti-VCAM1 mAb (clone M/K 2.7) (Bio X Cell) or rat IgG1 isotype control (clone HRPN) (Bio X Cell) every 3 days for 1 week.

### RNA isolation and bulk RNA-Sequencing

RNA was purified from MCMV-M45 tetramer^+^ CD8^+^T cells with the RNeasy Plus Micro Kit according to the manufacturer’s protocol (Qiagen). RNA was quantified with a Qubit flex fluorometer (Invitrogen) and the quality was assessed on a Bioanalyzer 2100 (Agilent) using a RNA 6000 Pico chip (Agilent). Samples with an RNA integrity number (RIN) of > 8 were used for cDNA amplification using Smart-Seq V4 Ultra Low Input RNA Kit (Takara). Libraries were made from 1ng of full-length cDNA using the Nextera-XT DNA Sample Preparation Kit (Illumina) according to the manufacturer’s instructions. The quantity of the cDNA libraries was assessed with Qubit flex fluorometer (Invitrogen) and the average library size was determined using Agilent’s 2100 Bioanalyzer HS DNA assay. Quantified libraries were sent to Novogene (Cambridge, UK) and sequenced on NovaSeq 6000 (Illumina) to generate around 30 mio PE150 reads for each sample.

### RNA sequencing analysis

The sequence reads were trimmed for adaptor sequences before being analyzed with Qiagen’s CLC Genomics Workbench (v23.0.2 with CLC’s default settings for RNA-Seq analysis). Raw RNA sequencing reads were mapped to the reference genome GRCm38.

### Single-cell RNA sequencing

Splenic CD11c^+^ cells were enriched with CD11c-MACS beads (Miltenyi Biotec), and then cells were labeled with the Mouse Immune Single-Cell Multiplexing Kit (BD Biosciences). Labeled cells have been counted and captured using the BD Rhapsody Single-Cell Analysis System Instrument (BD Biosciences) according to the provided instructions. Subsequently, whole transcriptome analysis (WTA) library preparation and Sample Tag library preparation were produced following the BD Rhapsody System mRNA WTA and Sample Tag Library Preparation Protocol. The quality and quantity of cDNA libraries were checked using the Agilent Bioanalyzer 2100 (Agilent), and sequencing (150 PE reads aiming for 50.000 raw reads/cell) was done at Novogene (Cambridge, UK) using the Illumina NovaSeq 6000 (Illumina). The generated raw data was preprocessed according to the Illumina standard protocol, and transcript alignment, counting, and demultiplexing were carried out utilizing the BD Rhapsody WTA analytic pipeline. Output matrices RSEC_MolsPerCell were further read into R (version 4.2.3) separately and converted to Seurat object using Seurat R package (*89*) (version 4.3.0.1). Genes with low expression level (detected in fewer than 3 cells) and cells that contained less than 200 detected genes were excluded from the analysis. Cells that did not meet the threshold of mean ± 3 times the median absolute deviation of mitochondrial gene number, total number of genes, and detected transcript number per cell for each separate sample were filtered out. Subsequently, the following processing and parameters were applied: normalization, variable feature search and scaling using sctranform, data integration, dimensional reduction by PCA and UMAP embedding (using 30 PCA dimensions), finding neighbors (using 30 PCA dimensions), and cluster search (resolution = 0.7). For each cluster, a main cell type was manually identified based on classical marker gene expression. The cluster identified as cDC1s was subjected to subsetting using “subset” function. Cell-cell communication analysis was performed using the CellChat R package. MMMs and Langerin^+^cDC1s were identified and subset from the mouse dataset using Seurat. Specifically, MMMs were defined as cells with Siglec1 expression > 0.1, while Langerin^+^cDC1s were defined as cells with CD207 expression > 0.1. These subsets were then used to create a CellChat object for downstream analysis, based on standardized procedures as previously described(*90, 91*). The inherent mouse database encompassing “secreted signaling’’, “ECM-Receptor’’ and “Cell-Cell contact’’ types of communication, was employed to infer intercellular communication networks based on known ligand-receptor interactions, facilitating the identification of key signaling pathways and interactions between these cell populations. Cell groups with fewer than 5 cells were filtered out using the filter Communication function in CellChat.

## Statistical Analysis

Data in all experiments are presented as mean ± SEM. Statistical analyses were assessed with two-tailed, unpaired Student’s t test to compare two groups or one-way ANOVA to compare multiple groups. Sample sizes were determined by experimental complexity, but no methods were used to determine normal distribution of the samples. Statistical analyses were performed using GraphPad Prism (version 9.4.0). The P value ≤ 0.05 was considered significant.

## Supplementary Materials

Fig. S1 to S6

## Acknowledgments

The authors would like to thank Dr. Khalad Karram, Nishada Ramphal, Elena Zurkowski, Zeynep Ergün, Theresa Schaller and Valeriya Zinina for their technical help. We are thankful to the Core Facility Flow Cytometry of the Research Center for Immunotherapy (FZI) for their support. We also thank Albert Nguyen and Lesley Ward for their technical help and for sending reagents. We are grateful to all members of the A.W. laboratory for their assistance and suggestions.

## Author contributions

X.Y.L. performed most experiments, analyzed data, and prepared the original draft of the manuscript. F.D. performed selected experiments and analyzed the data. M.A. and S.E. performed bioinformatics analyses of RNA sequencing data. M.K. supplied reagents and contributed sample preparation for RNA sequencing. D.B., A.E., M.B., A.E., K.C., L.J., D.U., and E.B. provide assistance with experiments and data analysis. H.C.P., J.L.G., B.E.C., and K.S.L. provided mice and discussion. N.H. provided intellectual inputs and suggestions. R.A., and J.d.H. provided reagents and reviewed the manuscript. A. S. provided technical support. T.B., E.v.S, and B.E.C. contributed to the review and editing of the manuscript. R.A.B. was involved in conceptualizing the study and reviewing and editing the manuscript. N.A.L. provided viral recombinants, helped with experiments and discussion. A.W. supervised the study and wrote the manuscript with X.Y.L.

## Data and materials availability

The accession numbers for the raw sequencing data reported in this paper will soon be available in a public repository. All other data needed to evaluate the conclusions in the paper are present in the paper or the Supplementary Materials.

## Supplementary Figures

**Fig. S1.**
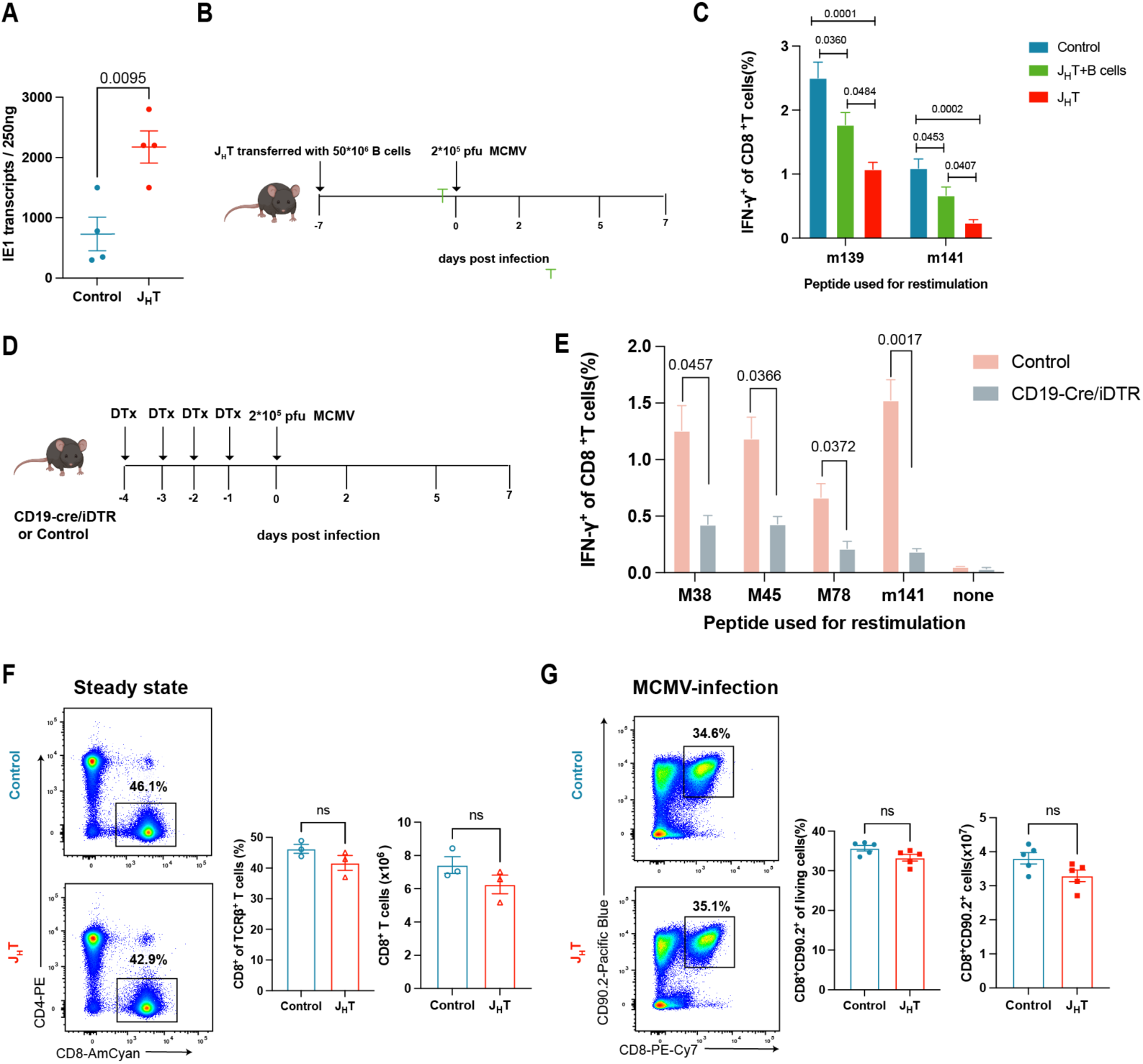
B cells are essential for the maintenance of the primary MCMV-specific CD8^+^ T cell response. (**A**) IE1 transcripts in the popliteal lymph nodes at 2 days post MCMV infection were measured by qRT-PCR (n = 4). (**B**) Schematic diagram outlining the program for reconstitution of J_H_T mice with WT splenic B cells; 1 week after B cell transfer, the mice were infected with MCMV. (**C**) Frequency of IFN-γ-producing CD8^+^ T cells in the spleens after stimulation with different dominant MCMV peptides at 7 days post-infection of control, J_H_T, or B-cell-reconstituted J_H_T mice (n = 7). (**D**) Schematic diagram outlining the program for CD19-Cre/iDTR and iDTR mice were treated with DTx according to the illustrated schedule, then the mice were infected with MCMV. (**E**) Frequency of IFN-γ-producing CD8^+^ T cells in the spleens after stimulation with MCMV peptides at 7 days post-infection of control or CD19-Cre/iDTR mice (n =3-7). (**F**)Representative flow cytometry dot plots of splenic CD8^+^T cells in control orJ_H_T mice at steady state. Bar graphs indicate the frequencies (left) and absolute numbers (right) of CD8^+^T cells in each group (n = 3). (**G**)Representative flow cytometry dot plots of splenic CD90.2^+^ CD8^+^T cells in MCMV infected control or J_H_T mice. Bar graphs indicate the frequencies (left) and absolute numbers (right) of CD8^+^T cells in each group (n = 5). Data are presented as the mean ± SEM and representative of two-three independent experiments. Statistical analysis: Student’s t-test, Fig. S1A, 1E,1F,1G. one-way ANOVA, Fig. S1C.

**Fig. S2.**
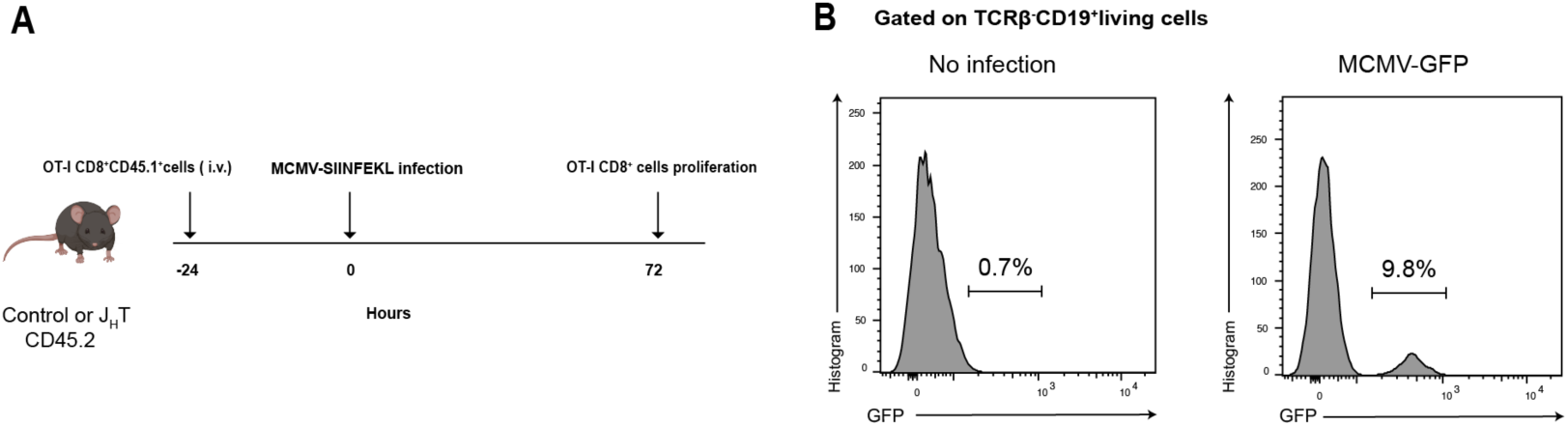
Loss of B cells reduces the priming of CD8^+^ T cells after viral infection (**A**) Schematic diagram outlining the program for OTI CD8^+^ T cells transfer experiments, 24 hours after OTI CD8^+^ T cell transfer, the mice were infected with MCMV-SIINFEKL. (**B**) Flow cytometry histograms of MCMV-EGFP^+^ B cells in the spleen of MCMV-EGFP infection WT mice and uninfected control WT mice.

**Fig. S3.**
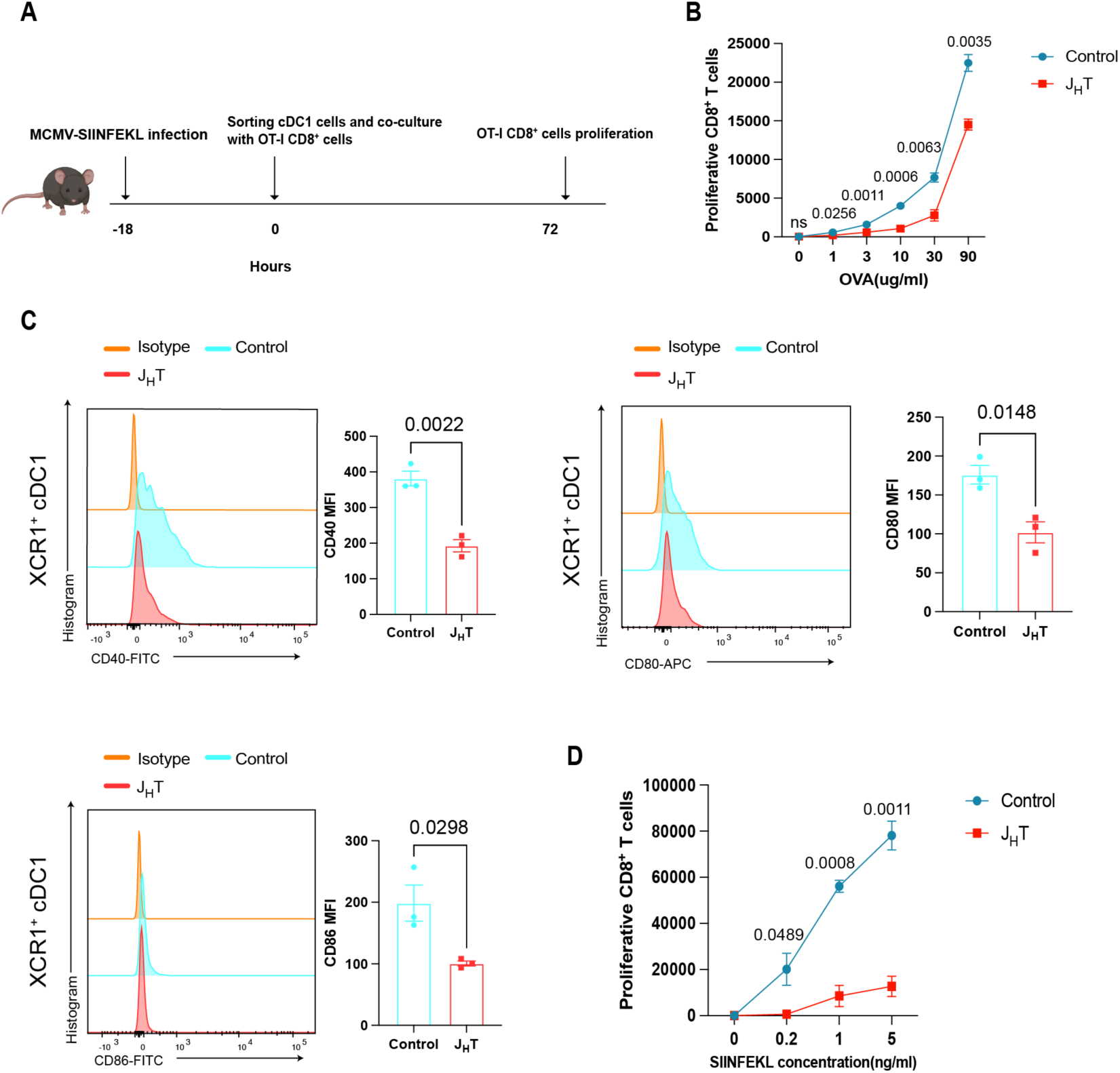
B cells contribute XCR1^+^CD8^+^cDC1s to prime CD8^+^ T cells. (**A**) Schematic diagram outlining the program used to co-culture OVA-specific transgenic OTI CD8^+^ T cells with cDC1s isolated from MCMV-SIINFEKL-infected control or J_H_T mice. (**B**) Line graph indicates the absolute numbers of proliferating OTI CD8^+^T cells co-cultured with cDC1s isolated from control or J_H_T mice and pulsed with OVA protein (n = 5). (**C**) Flow cytometry histograms showing the expression of CD40 (upper left), CD80 (upper right), CD86 (bottom left) in cDC1s from control or J_H_T mice. Bar graphs indicate the Median fluorescence intensity (MFI) of CD40 (upper left), CD80 (upper right), CD86 (bottom left) in cDC1s from control or J_H_T mice (n = 3). (**D**) Line graph indicates the absolute numbers of proliferating OTI CD8^+^T cells co-cultured with cDC1s isolated from control or J_H_T mice and pulsed with SIINFEKL peptide (n = 6). Data are presented as the mean ± SEM and representative of two-three independent experiments. Statistical analysis: Student’s t-test, Fig. S3B-S3D.

**Fig. S4.**
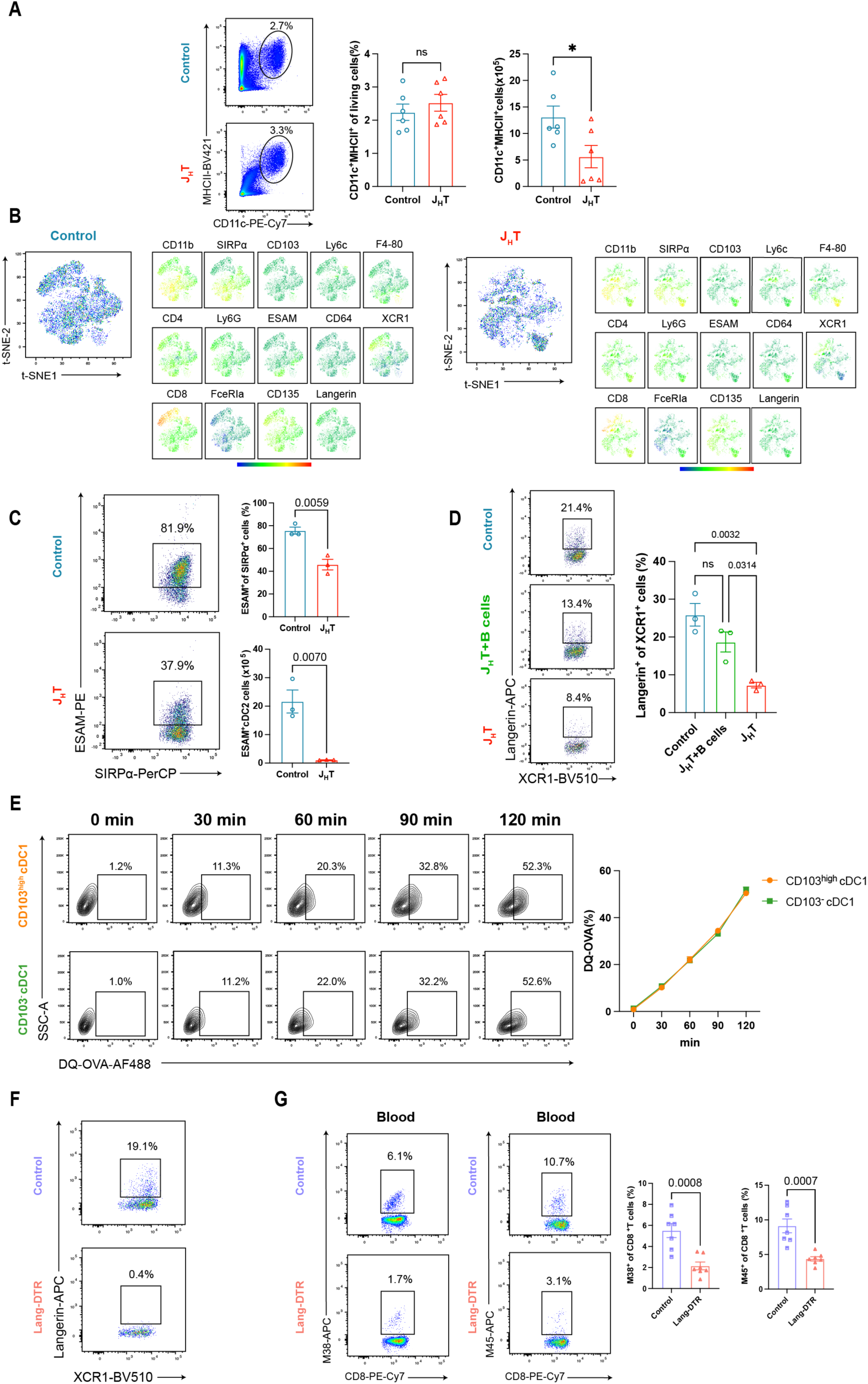
The homeostasis of splenic Langerin^+^ cDC1s is regulated by B cells. (**A**) Flow cytometry dot plots of CD11c^+^MHCII^+^ cDCs in the spleens of control or J_H_T mice. Bar graphs indicate the frequencies (left) and absolute numbers (right) of CD11c^+^MHCII^+^ cDCs (n = 6). (**B**) t-SNE of the expression of selected markers associated with the cDCs subgroup in the control (left) and J_H_T (right) cDCs populations (n = 3). (**C**) Flow cytometry plot showing splenic ESAM^+^ cDC2 cells in control or J_H_T mice. Bar graphs indicate the frequencies (top) and absolute numbers (bottom) of ESAM^+^ cDC2s (n = 3). (**D**) Representative flow cytometry plots of splenic Langerin^+^ cDC1s in control, J_H_T, or and B-cell-reconstituted J_H_T mice. Bar graphs indicate the frequencies of Langerin^+^ cDC1s (n = 3). (**E**) Flow cytometry plots of DQ-OVA^+^ XCR1^+^ cDC1s exposed to DQ-OVA-for different lengths of time. Briefly, CD103^+^cDC1s or CD103^-^cDC1s isolated from WT mice were incubated with 5 μg/ml OVA-DQ for 0, 30, 60, 90, or 120 minutes at 37 °C and 5% CO_2_. Line graph indicate the frequencies of DQ OVA^+^ XCR1^+^ cDC1s (n = 8). (**F**) Flow cytometry plot of splenic Langerin^+^ cDC1s in control and Lang-DTR mice after diphtheria toxin treatment. (**G**) Flow cytometry plot of MCMV-M38 (left) or MCMV-M45 (right) tetramer^+^ CD8^+^T cells in the blood of control or Lang-DTR mice. Bar graphs indicate the frequencies of MCMV-M38 ((left) or MCMV-M45 (right) tetramer^+^ CD8^+^T cells (n = 7). Data are presented as the mean ± SEM and representative of five (S4A and S4C) or two (S4B, S4D-S4G) independent experiments. Statistical analysis: Student’s t-test, Fig. S4A, S4C, S4E, and S4G. one-way ANOVA, Fig. S4D.

**Fig. S5.**
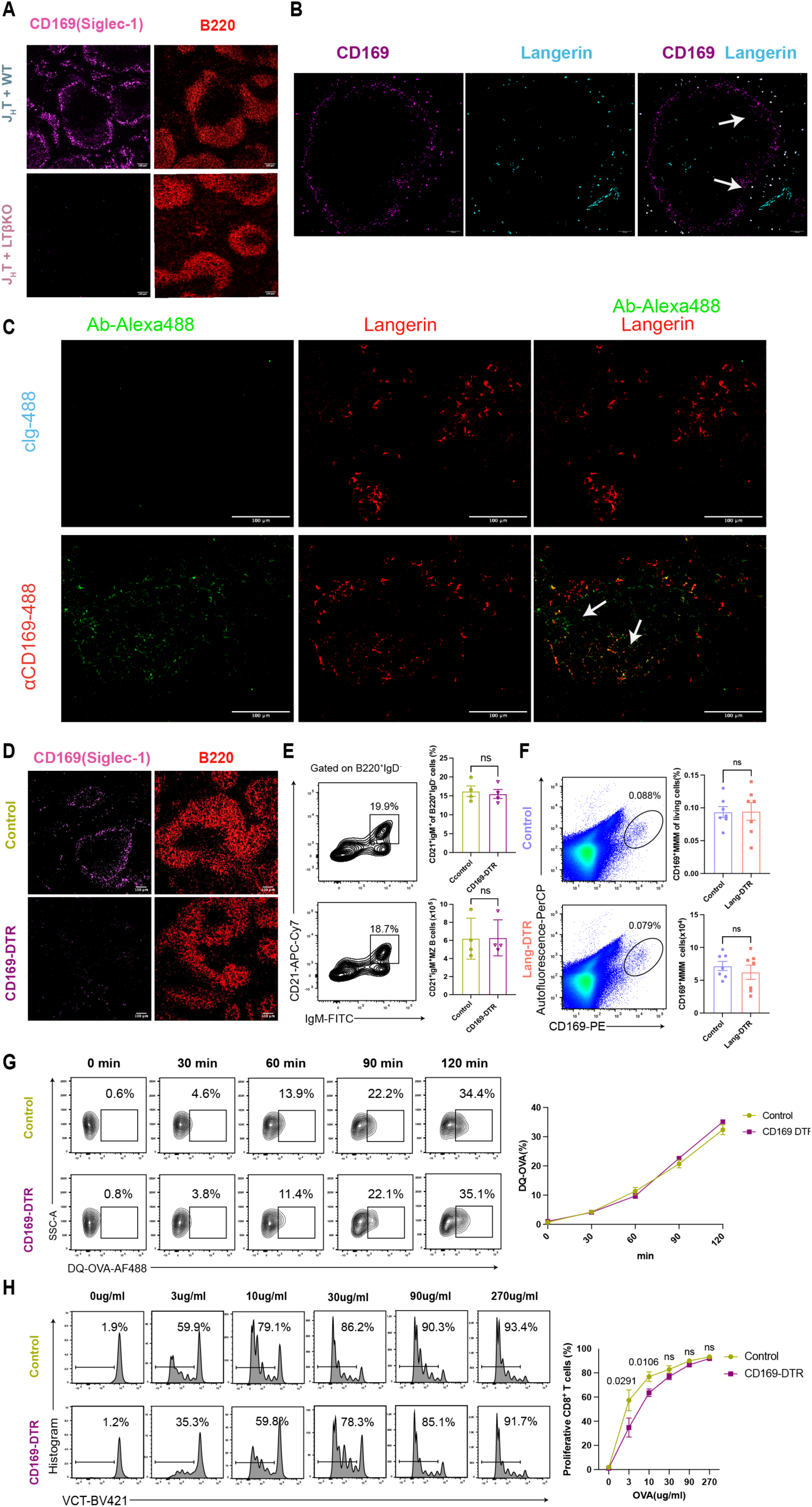
B cells expressing LTβ maintain MMMs to support homeostasis of splenic Langerin^+^ cDC1s (**A**) Immunofluorescence images of CD169 (Siglec-1) and B220 in the spleens of J_H_T mixed WT BM chimera mice and J_H_T mixed LTβ-deficient BM chimera mice (scale bar, 100 µm; n = 3). (**B**)Immunofluorescence images of CD169 (Siglec-1) and Langerin in the spleens of WT mice (scale bar, 50 µm; n = 4). (**C**) Immunofluorescence images of injected Ab-Alexa488 and Langerin in the spleens of WT mice (scale bar, 100 µm; n = 4) (**D**) Immunofluorescence images of CD169 (Siglec-1) and B220 in the spleens of control and CD169-DTR mice (scale bar, 100 µm; n = 4). (**E**) Flow cytometry contour plot of IgD^-^ CD21^+^ IgM^+^ MZ B cells in the spleens of control or CD169-DTR mice. Bar graphs indicate the frequencies (top) and absolute numbers (bottom) of MZ B cells (n = 4). (**F**) Flow cytometry dot plots showing MMMs in the spleens of control and Lang-DTR mice. MMMs were identified as live CD45^+^ autofluorescent CD169^+^ cells. Bar graphs indicate the frequencies (top) and absolute numbers (bottom) of MMMs (n = 6). (**G**) Flow cytometry plots of OVA-DQ^+^ XCR1^+^ cDC1s exposed to DQ-OVA for different lengths of time. Briefly, cDC1s isolated from control or CD169-DTR mice were incubated with 2 μg/ml DQ-OVA for 0, 30, 60, 90, or 120 minutes at 37 °C and 5% CO_2_. Line graph indicate the frequencies of DQ-OVA ^+^ XCR1^+^ cDC1s (n = 5). (**H**) Proliferation of OVA-specific transgenic OTI CD8^+^ T cells co-cultured with cDC1s isolated from control or CD169-DTR mice and pulsed with OVA protein. Line graphs indicate the frequencies of proliferating OTI CD8^+^T cells (n = 5). Data are presented as the mean ± SEM and representative of two independent experiments. Statistical analysis: Student’s t-test, Fig. S5E-S5H.

**Fig. S6.**
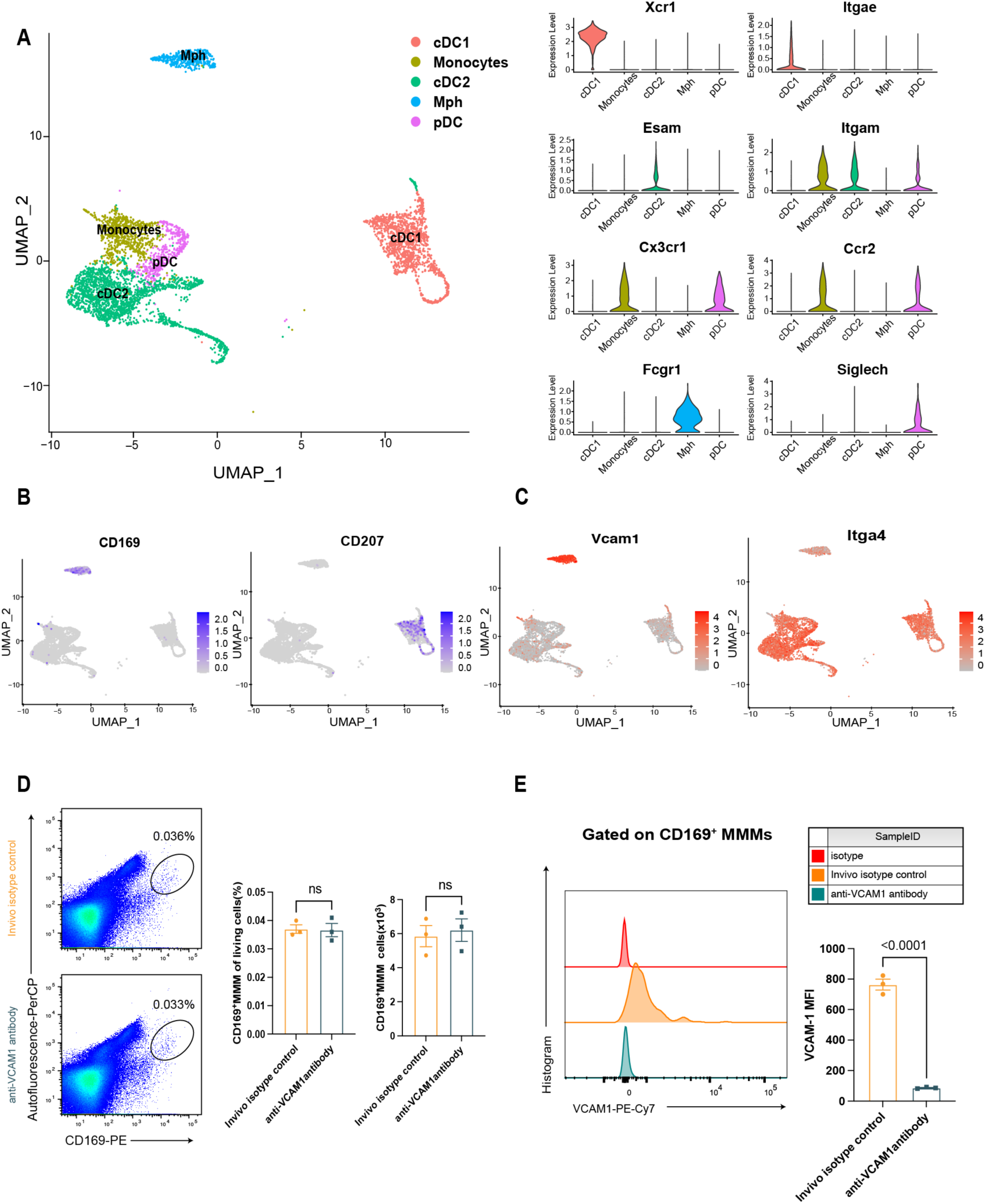
MMMs via VCAM1 - ITGA4/ITGB1 Cross-talk with Langerin^+^XCR1^+^cDC1s (A) UMAP plots of indicated clusters (left) and violin plots of their feature genes (right). (B) UMAP plots showing MMMs and Langerin^+^ cDC1s clusters. (**C**) UMAP plots showing VCAM1 and ITGA4 expression in indicated clusters. (**D**) Representative flow cytometry dot plots of MMMs in the spleens of control IgG or anti-VCAM1 antibody treatment mice. Bar graphs indicate the frequencies (left) and absolute numbers (right) of MMMs (n = 3). (**E**) Flow cytometry histograms showing the expression of VCAM1 in MMMs from control IgG or anti-VCAM1 antibody treatment mice. Bar graphs indicate the Median fluorescence intensity (MFI) of VCAM1from control IgG or anti-VCAM1 antibody treatment mice (n = 3). Data are presented as the mean ± SEM and representative of two independent experiments (S6D and S6E). Statistical analysis: Student’s t-test, Fig. S6D.

